# _Metabolic_ shift to serine pathway induced by lipids confers oncogenic properties in non-transformed breast cells

**DOI:** 10.1101/2024.02.21.581404

**Authors:** Mariana Bustamante Eduardo, Gannon Cottone, Curtis W. McCloskey, Shiyu Liu, Maria Paula Zappia, Elizaveta V. Benevolenskaya, A.B.M.M.K. Islam, Maxim V. Frolov, Flavio R. Palma, Peng Gao, Joel Setya, Hongyu Gao, Xiaoling Xuei, Rama Khokha, Jason Locasale, Marcelo G. Bonini, Navdeep S. Chandel, Seema A. Khan, Susan E. Clare

## Abstract

A lipid metabolism gene signature is associated with the risk of estrogen negative breast cancer (ER-BC). In vitro, lipid exposure alters histone methylation affecting gene expression and increasing flux through various metabolic reactions; but little is known about the mechanism(s) linking lipids and epigenetic reprogramming with the genesis of ER-BC. Here we show that the metabolism of the medium-chain fatty acid Octanoic Acid (OA) in preference to glucose and glutamine results in a metabolic shift toward the serine pathway increasing the production of SAM, glutathione, and 2-HG, with implications for oncogenesis: SAM production results in epigenetic fostered plasticity leading to reprogramming/selecting cells that express Neural, EMT and BC related genes. 2-HG exposure results in appearance of DNA breaks, potentially consequent to the inhibition of essential demethylases for HR repair. ROS increases shortly after OA exposure and is mitigated by antioxidant defenses, which favors/enables the survival of specific cell subtypes.

## Main

Breast cancer (BC)-related mortality is decreasing due to more effective therapy and to early detection. However, BC incidence continues to increase globally.^1^ This underscores the deficiency of current preventive strategies, particularly those that protect against estrogen receptor negative BC (ER-BC).^2^ To achieve subtype-specific cancer prevention, using agents that reverse, suppress or prevent carcinogenesis,^3^ an understanding of the etiology of the different subtypes of BC is needed. For example, premenopausal obesity, African American ancestry, *BRCA1* carrier status, and certain germline nucleotide variants are associated with a higher frequency of ER-BC.^4–7^ Nevertheless, the strongest risk factors for BC, other than germline mutations in tumor suppressor genes, are found in the breast, e.g., epithelial atypia and mammographic density.^8,9^

We have focused on examining the in-breast microenvironment to identify factors that promote ER-BC and that may be disrupted for prevention. Since studies of metachronous contralateral BC (CBC) show a similarity in the ER status of the CBC to the index primary,^10,11^ we began by studying the contralateral, unaffected breast (CUB) of patients with unilateral BC, using this as a model to discover potential markers of subtype-specific risk. We analyzed gene expression profiles of epithelial cells from CUBs and identified a lipid metabolism (LiMe) gene signature enriched in the CUBs of women with ER-BC.^12,13^ To better understand lipid metabolism in the breast, we studied the effect of fatty acids on non-transformed breast epithelial cells and observed increased flux through several metabolic reactions including those involved in serine, one carbon (1C) and glycine (SOG); methionine; and in reactions producing endogenous antioxidants. Moreover, lipid exposure resulted in profound changes in chromatin packing density, chromatin accessibility, histone posttranslational modifications and gene expression.^14^

Consulting metabolic reaction network reconstructions, the most likely hypothesis to account for the increase in SOG and methionine cycle metabolic flux is the shunting of metabolic activity from glycolysis to the de novo serine pathway. To determine how this occurs, its consequences and to further study the effects of fatty acids on the normal breast, we have performed metabolomics, epigenomic profiling and single cell RNA sequencing on cell lines and tissue-derived breast microstructures. We show that the metabolism of lipids in preference to glucose and glutamine re-wires cellular metabolism away from glycolysis and toward the SOG and methionine pathways increasing S-adenosylmethionine (SAM), glutathione and 2-hydroxyglutarate (2-HG) levels leading to survival, epigenetic fostered phenotypic plasticity and possibly metabolic BRCAness.

## Results

### Metabolism of fatty acids in preference to glucose and glutamine results in a metabolic shift toward the de novo serine pathway

The metabolic flux analysis of bulk RNA-seq data that we previously published revealed that Octanoic Acid (OA) drives flux through many metabolic reactions including those involved in SOG and methione pathways.^14^ To test whether in presence of OA, carbons from glucose are being diverted into the SOG and methionine pathways therby increasing the production of the main methyl donor SAM; we performed ^13^C-glucose tracing in MCF-10A cells exposed to OA. Twenty-four-hour exposure to OA resulted in increased fractional abundance of SAM M+1 through the serine pathway (Fig. 1a); in addition, it decreased phosphoribosyl pyrophosphate (PRPP) M+5 and fractional abundance of ATP M+5-7 (Extended Data Fig.1a). ^13^C metabolic flux analysis revealed that upon OA exposure, glucose uptake, glycolysis and pentose phosphate pathway flux decreased (Extended Data Fig.1b); and 1C-THF was redirected to the methionine cycle increasing flux to DNA methylation (Fig.1b). OA treated cells had an increased Cancer index (defined as the lactate ratio) (Fig 1b), Non-canonical TCA index and TCA-index (Extended Data Fig.1c).

**Fig. 1:**
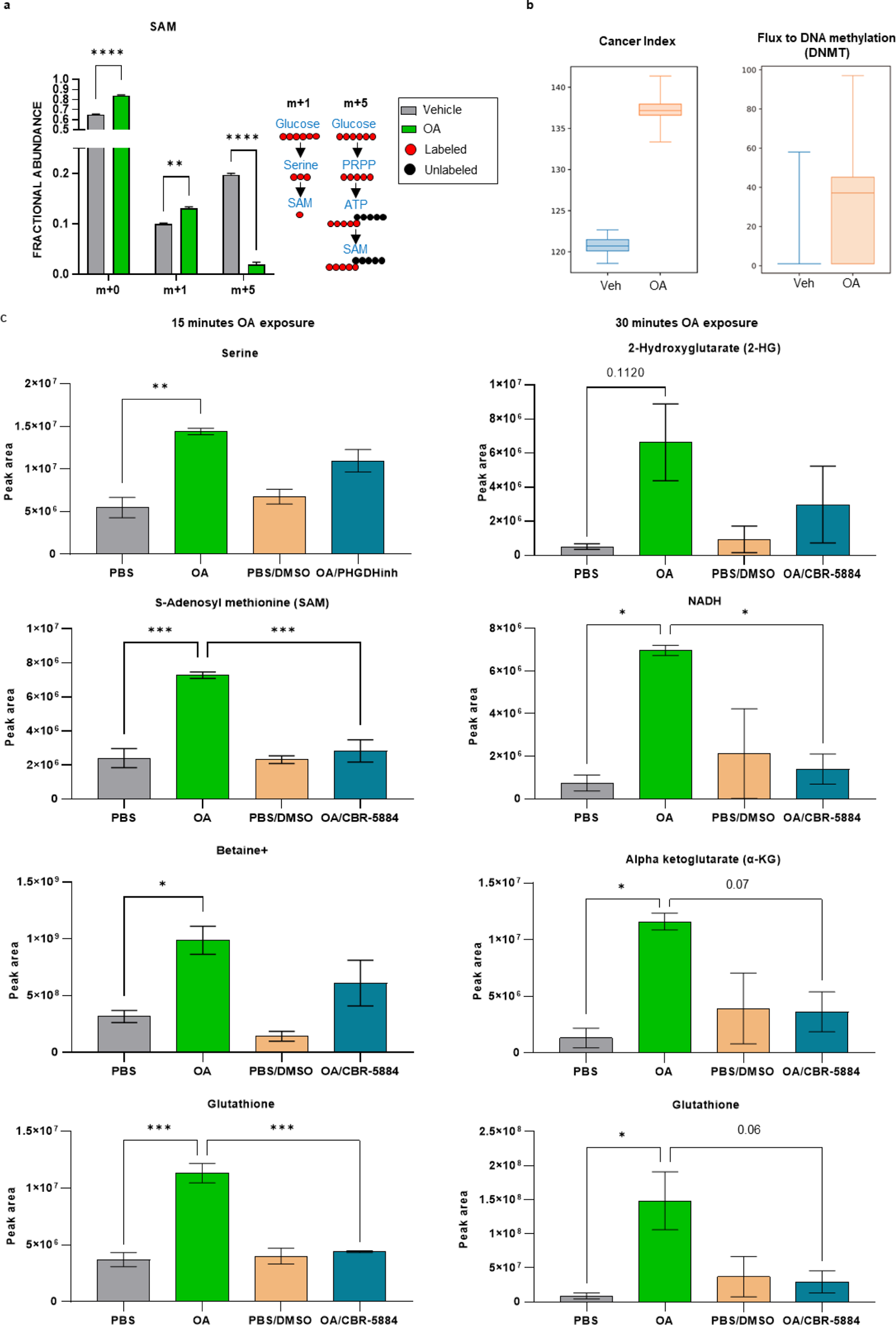
Metabolism of Octanoic Acid (OA) results in a metabolic shift toward the de novo serine pathway. **a,** Fractional abundance of S-adenosyl methionine (SAM) isotopologues. *p < 0.05, ****p < 0.0001 (2way anova with Turkey test, n=3, mean with sd). 4 hours labeling. On the right schematic of derivation and contribution of carbon atoms in SAM synthesis. **b,** 13C Metabolic flux calculations. Cancer Index was calculated as the lactate ratio. **c,** Measument of serine, S-adenosyl methionine (SAM), Betaine+, Glutathione, 2-Hydroxyglutarate (2-HG), NADH and alpha ketoglutarate (αKG) after 15-and 30-minutes exposure to PBS (vehicle), OA, PBS plus DMSO (vehicle) and OA plus CBR5884. *p < 0.05, **p < 0.01, ***p < 0.001 (2way anova with Turkey test, n=3, mean with SEM).

We hypothesized that the first and rate limiting step in the *de novo* serine pathway, which is catalyzed by Phosphoglycerate Dehydrogenase (PHGDH) is key to these observations. In the forward direction, PHGDH participates in SOG and methionine pathways that produce SAM and in the reverse direction produces the oncometabolite 2-HG. We therefore performed targeted metabolomics in MCF-10A cells exposed to OA in presence or absence of the PHGDH inhibitor CBR-5884. This revealed the increased production of Serine, SAM, glutathione and Betaine after 15 minutes OA exposure; and the increased production of 2-HG, glutathione, NADH and alpha ketoglutarate (αKG) after 30 minutes OA exposure (Fig. 1c). Blocking PHGDH with CBR-5884 prevented these increases.

These results show that the metabolism of lipids in preference to glucose and glutamine re-wires cellular metabolism away from glycolysis and toward the SOG and methionine pathways increasing the production of endogenous antioxidants such as glutathione, the main methyl donor SAM and the oncometabolite 2-HG.

### Cells deploy additional antioxidant defenses to control the concentration of ROS

Upon OA exposure cells deploy additional antioxidant defenses such as glutathione, NADH and αKG (Fig. 1c). In addition, previous metabolic flux analysis from bulk RNA-seq data also showed an increase of mitochondrial catalase and ALDH1L1.^14^ We postulated that increased flux in the methionine cycle would lead to incresed transsulfuration pathway activity leading to increased production of glutathione to mitigate the effects of increased reactive oxygen species (ROS) due to fatty acid oxidation. Therefore, we monitored ROS-induced redox changes live for 60 minutes in MCF-10A cells transduced with the ORP1-roGFP2 vectors with either mitochondrial or nuclear localization. This revealed a significant increase of mitochondrial (Fig. 2a) and nuclear (Fig. 2b) ROS after 5 minutes OA exposure. Mitochondrial and nuclear ROS levels remained significantly high after 60 minutes of exposure. Mitochondrial and nuclear ROS significantly peaked (p < 0.0001) after 15 minutes with OA and slowly decreased. Nuclear ROS decreased much more slowly than mitochondrial ROS. Thus, OA exposure results in the deployment of additional antioxidant defenses to mitigate increased concentrations of ROS.

**Fig. 2:**
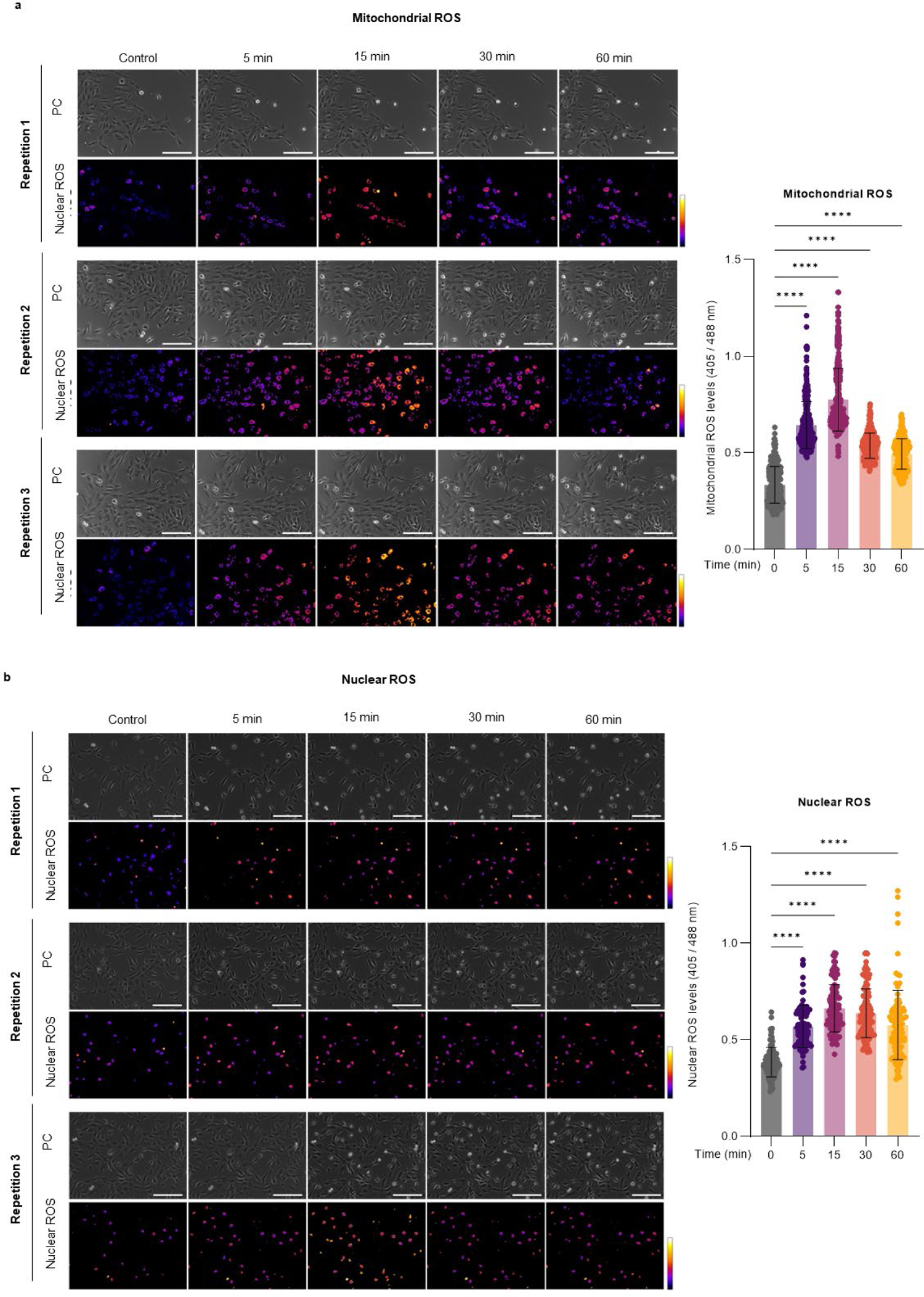
Reactive oxygen species (ROS) life monitoring in MCF-10A cells. **a**, mito-roGFP2-Orp1 monitors mitochondrial redox state in MCF-10A cells. Ratio of emission at 405 and 488 nm obtained as a response of OA exposure. Scale bar 200µM. **b**, nls-roGFP2-Orp1 monitors nuclear redox state in MCF-10A cells. Ratio of emission at 405 and 488 nm obtained as a response of OA exposure. Scale bar 200µM., ****p < 0.0001 (Ordinary one-way ANOVA with Turkey test, mean with SD).

### The increase of SAM and 2-HG leads to histone methylation affecting gene expression

We postulated that the increase of the main methyl donor SAM leads to the increased histone methylation we observed in our earlier publication;^14^ additionally, the increase of 2-HG, acting as an inhibitor of α-KG-dependent dioxygenases such as histone demethylases,^15^ would contribute to the increased methylation. We posited that H3K4me3, a histone mark associated with transcriptionally active chromatin, and H3K27me3, a histone mark associated with repression of gene expression, explain the profound gene expression changes caused by OA.^14^. CUT&RUN for H3K27me3, revealed twelve peaks differentially enriched in control (FDR < 0.05) and associated with increased gene expression in OA (Extended Data Fig.2a-b, Supplementary table 1a). Among them were stem cell markers LGR6 (upregulated by OA, log2FC = 1.9)^14^, a stem cell marker also associated with ER-BC, and PLAG1 (upregulated by OA, log2FC = 2.8)^14^. Interestingly with regard to the increase in LRG6 expression, the percentage of ALDH+ cells, a marker of stem cells, increased by at least 10% after OA exposure (Extended Data Fig.2c). H3K4me3 CUT&RUN identified 661 differential peaks significantly enriched in OA treated MCF-10A cells (Supplementary table 1b). 73% of H3K4me3 OA-associated peaks were in regulatory regions of OA-induced genes (FDR < 0.01)^14^ (Fig.3a). Homer motif analysis revealed an overrepresentation of binding sites for transcription factors (p < 0.05) associated with EMT (Zeb1, Slug), neural functions (E2A, AP1, JunB), neuronal-injury (Atf3), serine pathway (Atf3, Atf4) and stress response (Chop, Atf3, Atf4) (Fig.3b). Examples of significantly upregulated genes upon OA with significant H3K4me3 enrichment are shown in Fig. 3c. We used Enrichr [https://maayanlab.cloud/Enrichr/] to perform pathway analysis of OA-induced genes with increased H3K4me3 peaks. Among the overrepresented pathways were those involved in EMT, neural related pathways, embryonic stem cell pluripotency, cell migration and invasion, and breast cancer (Fig.3d). To determine if inhibition of enzymatic reactions in the *de novo* serine pathway and/or H3K4 trimethylation block the OA-induced changes in gene expression, MCF-10A cells were exposed to OA in presence or absence of the PHGDH inhibitor CBR-5884 or the histone methyltransferase inhibitor Piribedil. Gene expression of OA-induced with and without H3K4me3 peaks was then measured by real-time quantitative PCR (rt-qPCR) (Fig.3e). Blocking PHGDH significantly prevented the induction of the OA-induced stress-response gene ATF4; the induction of ATF3 and CHOP was higher in presence of the inhibitor and the expression of PERK did not change. Piribedil significantly induced the expression of ATF3, ATF4 and CHOP without affecting PERK. OA induced the expression of PHGDH which was prevented by CBR-5884. NANOG induction was prevented by both CBR-5884 and Piribedil. The expression of H3K4me3 controlled genes NGFR, PARM1, AGR2 and PDGFRA were prevented by blocking the de novo serine pathway and histone methylation. The increase of NGF was prevented by CBR-5884 while Piribedil induced its expression further. Overall, these results show that many gene expression changes driven by the metabolic flip towards the SOG and the methionine cycles occur as a result of epigenetic reprogramming secondary to the increase of H3K4me3.

**Fig. 3:**
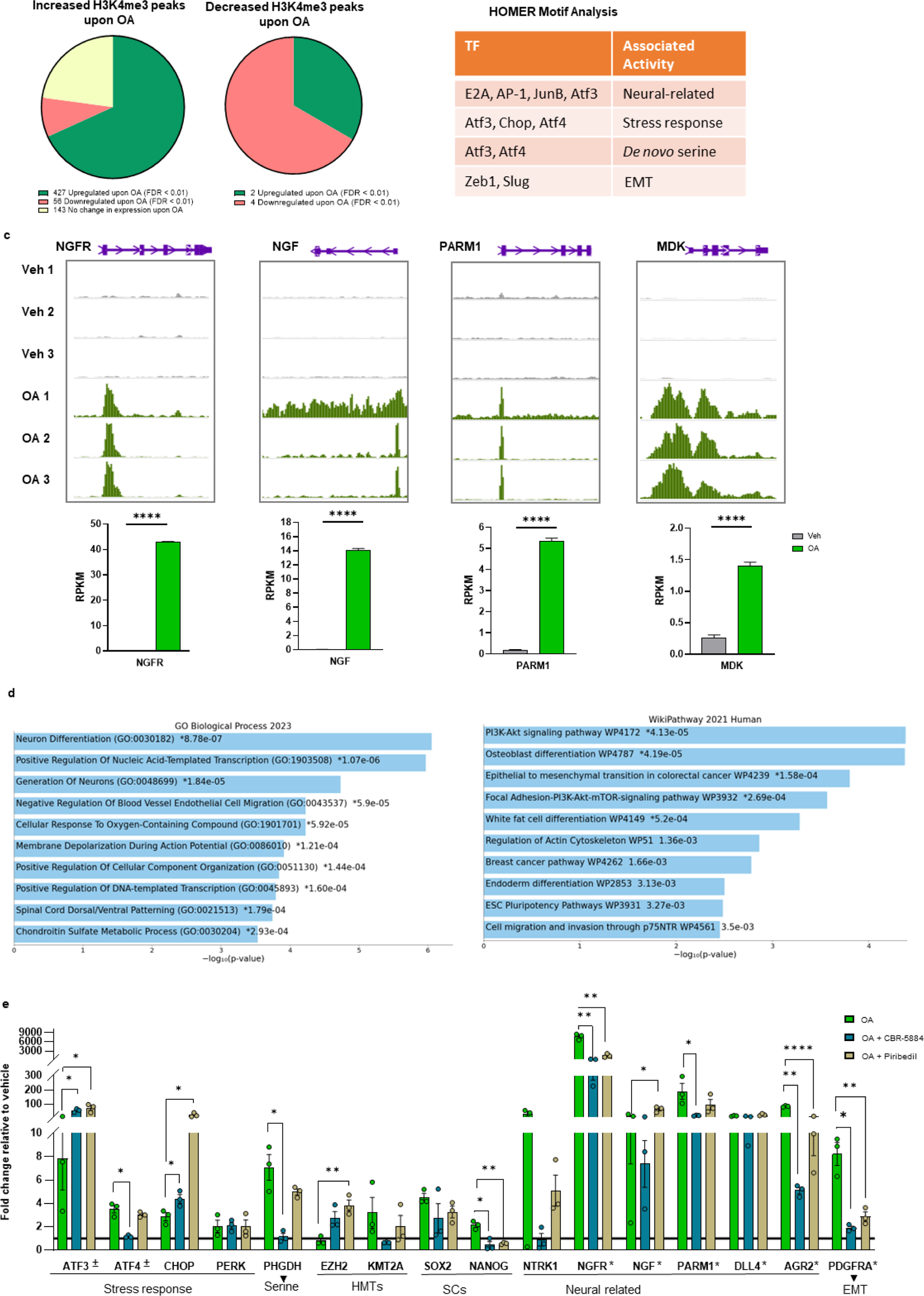
CUT&RUN for H3K4me3 upon OA exposure. **a,** Pie chart depicting the distribution of OA-associated H3K4me3 peaks (FDR < 0.01) at OA-modulated genes (FDR < 0.01). **b,** Table depicting relevants transcription factor (TF) motifs identified with HOMER in increased H3K4me3 peaks and their associated activity. **c,** Histograms showing H3K4me3 occupancy at NGFR, NGF, PARM1 and MDK in vehicle (grey tracks) and OA (green tracks). The peaks are visualized with the WashU Epigenome Browser. The lower panel shows boxplots of RNA expression RPKM values. ****p < 0.0001. **d,** Gene Ontology (GO) Biological Process 2021 and WikiPathway 2021 classifications for genes overexpressed upon OA treatment and associated with increased H3K4me3 peaks. **E,** Boxplots showing the expression of OA-induced genes upon OA, OA plus CBR-5884 and OA plus Piribedil measured by qPCR. *OA-induced genes with enriched H3K4me3 peaks, ^±^ Promotes the transcription of *de novo* serine genes. *p < 0.05, **p < 0.01, ****p < 0.0001 (Multiple unpaired t test, mean with SEM).

### OA-exposed MCF-10A cells adopt a neural-like phenotype when grown on Poly-D-Lysine/Laminin coated plates

Given the strong signal of neural differentiation noted above (Fig. 3d), which we had also observed in the bulk RNA-seq data^14^, we sought to determine if OA-exposed MCF-10A cells adopt a neural phenotype in cell culture. Poly-D-Lysine and laminin are utilized in neural cell culture to promote cell attachment, growth, and differentiation. Therefore, OA-exposed MCF-10 cells were cultured on Poly-D-Lysine/Laminin coated plates (Extended Data Fig.3). Culture of MCF-10A in presence of OA clearly results in the switch to a neural phenotype complete with the outgrowth of neurites and a cell body that is polygonal.

### OA exposure leads to DNA breaks

OA exposure leads to the significant increase of D-2-HG (Fig.4a) but not L-2-HG (Fig.4b). D-2-HG, produced by PHGDH, was no longer induced when PHGDH was silenced. Among the αKG-dependent dioxygenases inhibited by 2-HG are KDM 4A/B which are required for homologous recombination (HR) repair.^16^ Therefore, we performed an alkaline comet assay to determine whether OA exposure leads to DNA breaks. This revealed that OA significantly increased DNA breaks in MCF-10A cells. Alkaline comet assay showed that OA and exogenous 2-HG significantly increased comet tails. However, using the PHGDH inhibitors CBR5884 or NTC-503 did not decrease comet tails. This is likely a consequence of increased ROS due to the reduction of glutathione upon PHGDH inhibition. Significantly increased comet tails were also observed in MCF-10A cells expressing doxycycline-induced PHGDH likely due to the overexpression of this gene which leads to the increase of 2-HG (Extended Data Fig.4). It is likely that OA-induced increase of 2-HG contributes to the inhibition of the αKG-dependent dioxygenases required for HR repair resulting in DNA damage.

### OA affects the cellular composition of breast microstructures

To explore the effects of lipids in breast tissue ex vivo; we exposed breast microstructures derived from reduction mammoplasty tissue from 5 donors to vehicle or OA and measured OA-induced genes previously reported^14^ by real-time quantitative PCR (rt-qPCR). Most of the selected genes were induced by OA in tissue derived microstructures (Extended Data Fig. 5a). Because breast microstructures retain the original architectural integrity as well as the different cell subtypes of the breast environment;^17^ we further explored the effects of OA by single cell RNA sequencing (scRNA-seq). For this, breast microstructures from 2 donors were exposed to vehicle and OA for 24 hours, dissociated into single cells and approximately 10,000 individual cells per sample were sequenced using the 10x Genomics platform. After quality control and adjusting for technical noise, single-cell transcriptomic profiles for 36,904 cells and 41,445 genes were selected for the analysis (see “Methods” section). We performed Uniform Manifold Approximation and Projection (UMAP) with a resolution of 0.8 and a total of 25 clusters were identified including epithelial, fibroblasts, endothelial cells, and immune cells (Fig.5a). Cell populations were identified using the markers provided in Reed et al.^18^ Within the epithelial compartment, we have identified several subclusters of luminal progenitor (LP, 1-5), hormone sensing (HS, 1-4), basal (BSL, 1 and 2) cells and cells described by Reed et al. (29) as well as donor derived clusters (DDC, 1 and 2). OA greatly affected the proportion of many cell subtypes. Notably, within the epithelial compartment the proportion of BSL1, HS1 and LP3 increased by OA and the percentage of DDC2, HS2, LP1 and LP4 decreased. In addition, the proportion of EC angiogenic tip cells decreased and the proportion of EC venous increased upon OA.

### OA modulates gene expression in tissue-derived breast microstructures

OA modulated gene expression within each subcluster, Fig.5b displays the previously identified OA responsive genes (ATF3, PHGDH, NGF, NGFR, PARM1). About 58% of OA-induced genes in breast microstructures overlapped with OA induced genes in MCF-10A cells,^14^ and about 4% of them had H3K4me3 peaks (Fig. 6a). The expression of many genes within each subcluster was modulated by OA. Notably, in BSL1, LP3 and HS1, OA induced the expression of genes involved in lipid droplets, EGFR signaling, serine, glycine, and folate as well as endoplasmic reticulum stress, oxidative stress, neural and cancer related genes while lineage markers were repressed (Supplementary table 2). Notably, Reactome pathway analysis revealed an upregulation of terms related to neuronal system, signal transduction, DNA repair and metabolism, and a downregulation of terms related to cell-cell communication and extracellular matrix organization upon OA exposure (Supplementary table 3).

**Fig. 4:**
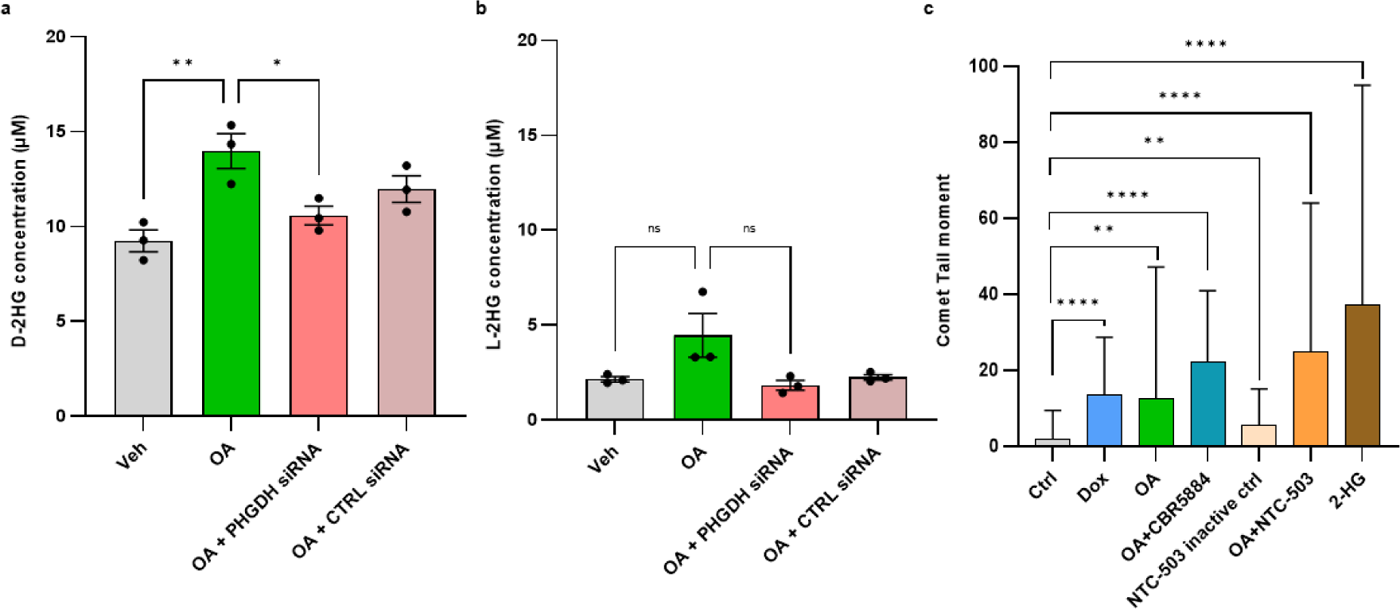
2-HG and DNA damage. **a,** Quantification of D-2-HG. *p < 0.05, **p < 0.01 (2way ANOVA with Turkey test, n=3, mean with SEM). **b,** Quantification of L-2-HG. ns = not significant (2way ANOVA with Turkey test, n=3, mean with SEM). **c,** Comet tail moment for control MCF-10A (ctrl), PHGDH-overexpressing MCF-10A (Dox), MCF-10A in presence of OA ± PHGDH inhibitors or inhibitor control and 2-HG. 2way ANOVA with Turkey test, mean with SEM.

**Fig. 5:**
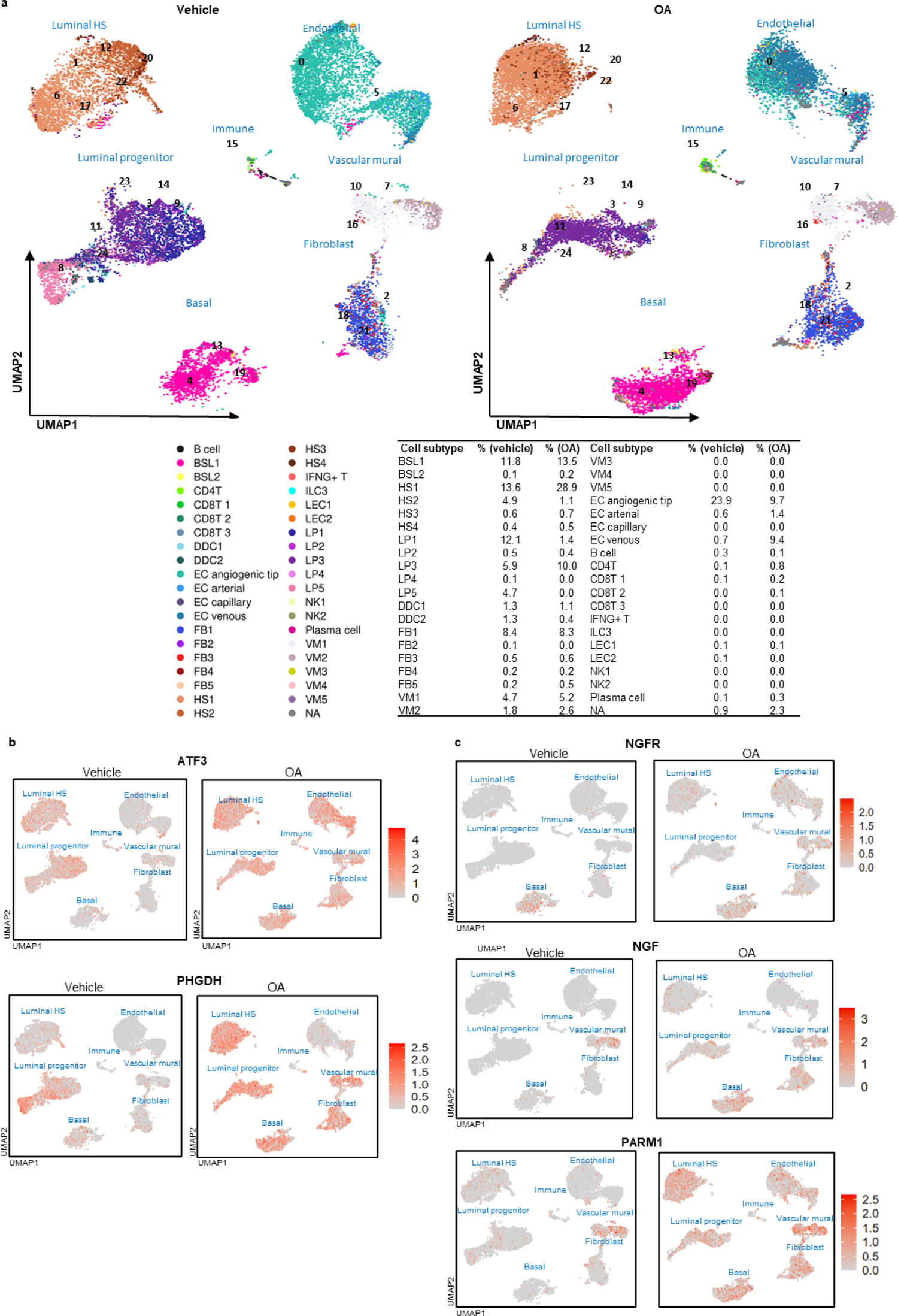
Single-cell landscape of vehicle and octanoic acid (OA) treated tissue-derived breast microstructures. **a**, Approximation and Projection (UMAP) plot of 36’904 cells identified a total of 25 different cell clusters. Cell subtypes defined by Reed et al. Table shows cell subtypes proportions. **b**, Feature plot of cells that are labeled according to *ATF3* and *PHGDH* transcription expression. **c,** Feature plot of cells that are labeled according to neural-related genes *NGFR*, *NGF* and *PARM1* transcription expression.

**Fig. 6:**
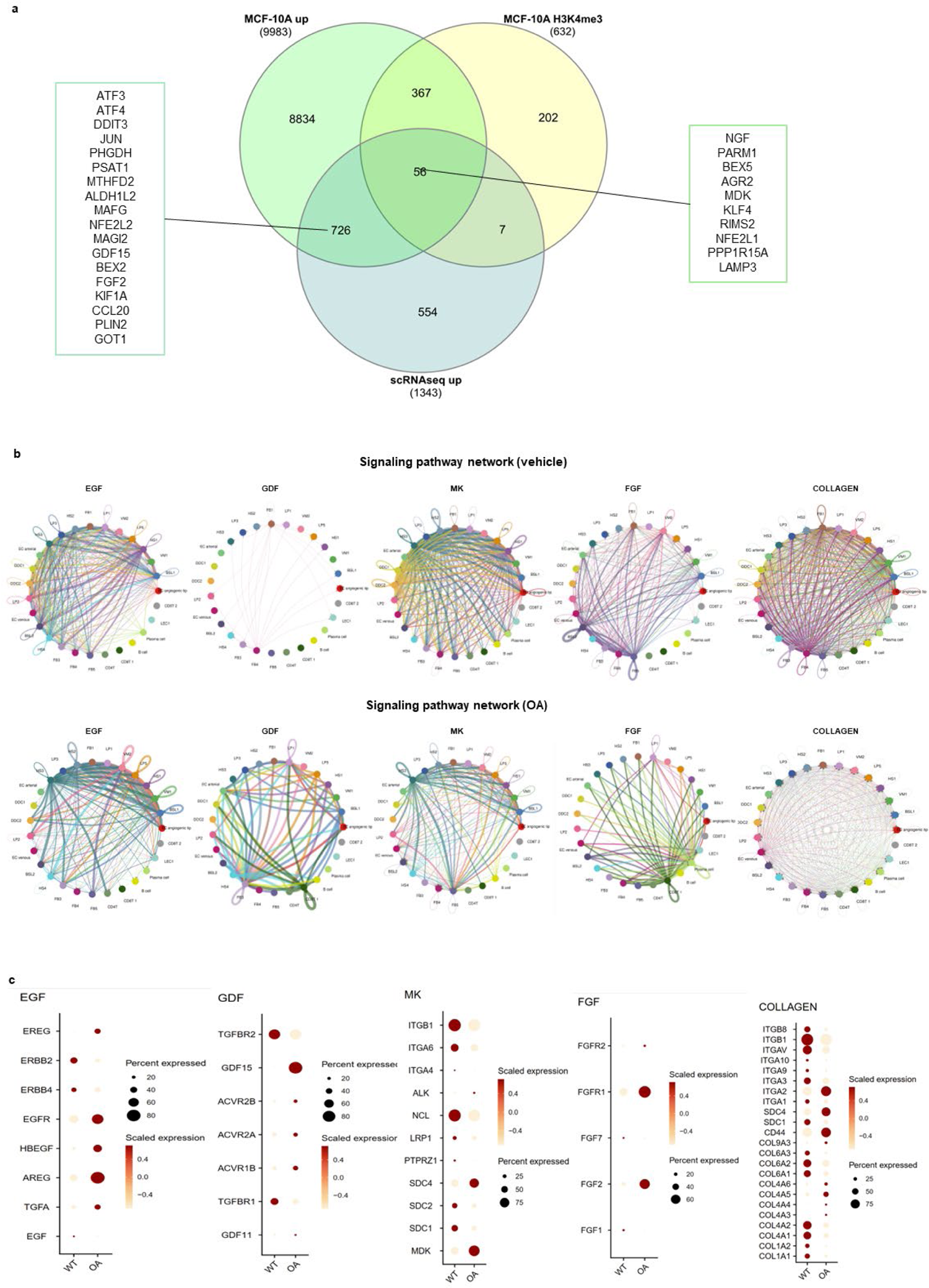
Single-cell RNA sequencing (scRNA-seq) analysis. **a**, Venn diagram depicting genes upregulated by OA in MCF-10A, genes with increased H3K4me3 peaks upon OA in MCF-10A and upregulated genes by OA in breast microstructures measured by scRNA-seq. **b,** Circle plot of a selection of signaling network in OA and vehicle **c,** Dot plot showing the expression of genes from a selection of signaling pathway in vehicle and OA.

OA modulates gene expression within each cell subtype leading to the downregulation of lineage markers, notably in BSL1 and LP3. Therefore, we wondered whether OA induced changes in epithelial cell state. To answer this question, we remapped the cell populations using epithelial cell state markers provided in Kumar et al.^19^ (Extended Data Fig. 5b). Kumar and colleagues have identified three epithelial compartments, the luminal hormone-responsive (LumHR), luminal secretory (LumSec) also known as LP and basal-myoepithelial (basal); and within each compartment they have described several cell states. Extended Data Fig. 5b shows the proportion of cell states in vehicle and OA. Upon OA basal cells slightly increased from 10% to 11%. Major changes were observed in the lumHR and lumsec compartments. Within the lumHR compartment most cells are in the lumHR-active state in the vehicle condition while upon OA the proportion of lumHR-active decreased (from 13% to 6%); the proportion of lumHR-SCGB slightly increased (from 2% to 3%); and the proportion of lumHR-major augmented from about 4% to 20%. Within the lumSec compartment, the main OA-induced changes were observed in lumSec-basal (decreased from 19% to 8%); lumSec-prol (decreased from 5% to 0%); and lumSec-HLA (augmented from 1% to about 9%).

This analysis shows that OA changed cell subtype proportions and clearly modulated gene expression within each cell compartment probably leading to a change in cell state or eventually a change in cell subtype.

### Cell-cell communication landscape changes upon OA exposure

We used CellChat to explore the effect of OA in cell-cell communications. Enriched ligand-receptors pairs with upregulated ligands in OA were different from those in vehicle. Among the signaling pathways that were highly active in OA were Epidermal Growth Factor (EGF), Growth Differentiation Factor (GDF), Midikine (MK), fibroblast growth factor (FGF) while Collagen was among the highly active in vehicle (Fig. 6b).

The three epithelial compartments that showed an increase proportion upon OA and changes of gene expression were BSL1, LP3 and HS1. The strongest signals from BSL1 to their specific receptors in most epithelial, stromal, and immune cells upon OA were AREG (EGF signaling ligand), GDF15 (GDF signaling ligand), MDK (MK signaling ligand), ANGPTL4 (ANGPTL signaling ligand), PTPRM (PTPRM signaling ligand) and NAMPT (VISFATIN signaling ligand) (Extended Data Fig. 6). The LAMININ and COLLAGEN pathways were among the strongest contributors in cell-cell communication in vehicle (Extended Data Fig. 7). Similarly, the strongest signals from LP3 and HS1 were AREG, GDF15, MDK, ANGPTL4, PTPRM and NAMPT to their specific receptors in most epithelial, stromal and immune cells in OA (Extended Data Fig. 6). The strongest signals from LP3 to most of the cells in vehicle were from MIF, CDH1 and APP pathways (Extended Data Fig. 7). In HS1, the strongest signals to most of the cells were from APP, CDH1, LAMININ and COLLAGEN pathways in vehicle. AREG, MDK, NAMPT and GDF15 signals from HS1 to some cell subtypes were also observed in OA (Extended Data Fig. 6). These results show that the overall interaction between cell subtypes changed upon exposure to OA, suggesting an increase of secreted signaling, a decrease of extracellular matrix-cell interactions and a decrease of cell-cell adhesions.

### Compass algorithm predicts metabolic systems affected by OA in BSL1, LP3 and HS1 cell populations

We performed metabolic flux analysis using our scRNA-seq data using Compass. Compass predicted differentially active metabolic reactions between OA and vehicle treatment in BSL1, LP3 and HS1 cells. This analysis showed that OA affects differently these three epithelial compartments; while greater OA effects are observed in BSL1 and HS1, lesser OA effects are observed in LP3. Compass predicted increased dependence of BSL1, LP3 and HS1 cells on Glycine, serine, alanine and threonine metabolism upon OA exposure (Fig. 7a); notably, key players in serine metabolism such as PHGDH, PSAT1 and Phosphoserine Phosphatase (PSPH) were enhanced by OA in these cell subtypes. Compass highlighted differences in ROS detoxification and glutathione metabolism among BSL1, LP3 and HS1. Catalase and superoxide dismutase reactions were enhanced by OA in BSL1. In BSL1 and HS1 cells, glutathione metabolism pathway’ reactions were increased in vehicle and OA; but notably, glutathione peroxidase (mitochondria) and glutathione NAD+ oxidoreductase were increased only in OA (Fig. 7b). In LP3 cells, glutathione metabolism pathway was enhanced in vehicle. The algorithm also predicted increased metabolic activity in tyrosine metabolism in all three cell subtypes (Fig. 7c). Remarkably, metabolic reactions that result in the synthesis of neurotransmitters were enhanced in BSL1, LP3 and HS1 (Fig. 7c). For example, dopamine beta-monooxygenase and tyrosine 3-monooxygenase in all three subtypes; Dopamine Sulfotransferase, Norepinephrine Sulfotransferase and noradrenaline N-methyltransferase in BSL1 and HS1. This analysis confirmed the upregulation of several metabolic reactions by OA; notably it enhanced the serine pathway in BSL1, LP3 and HS1; increased ROS detoxification and glutathione metabolism in BSL1; and it led to the synthesis of neurotransmitters in all three subtypes.

**Fig. 7:**
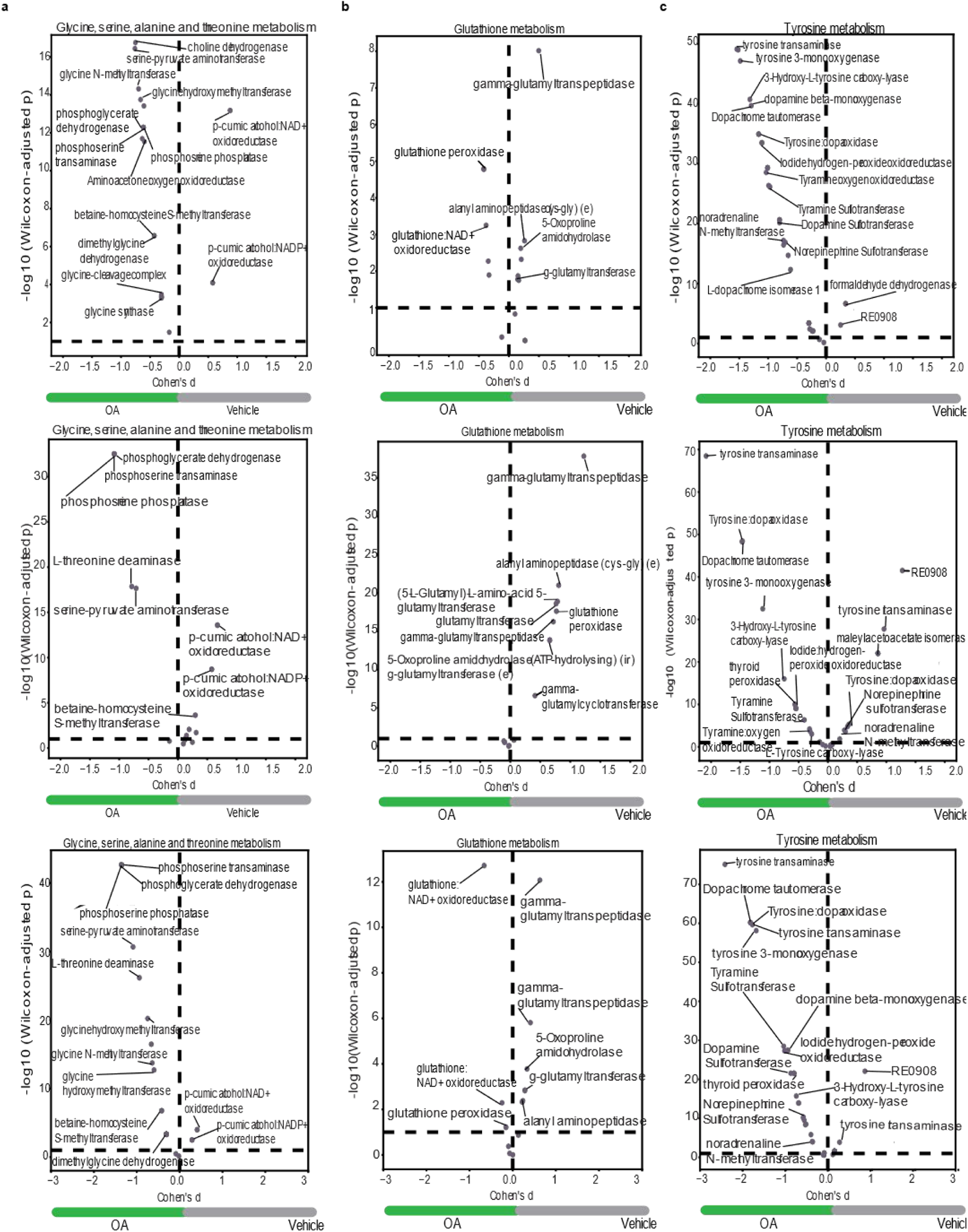
Octanoic acid (OA) effect in non-transformed breast cells. Specific reactions in BSL1 (upper panel), LP3 (middle panel) and HS1 (bottom panel) of Glycine, serine, alanine and threonine metabolism **(a),** Glutathione metabolism **(b),** and Tyrosine metabolism **(c). C**omplete list of reaction in supplementary tables 8-10.

## Discussion

Previous work from our group has focused on defining the in-breast environment that facilitates the development of ER-BC. We have identified a LiMe gene expression signature enriched in the CUBs of women with ER-BC.^12,13^ Further studies performed to discover the mechanism, i.e., the biologic basis linking lipid metabolism with the genesis of ER-BC, revealed that exposure of non-transformed breast epithelial cells to OA greatly affects gene expression, chromatin architecture, post-translational histone protein modifications and metabolic flux.^14^

We began these experiments using 5 mM OA and MCF-10A cells. OA, for the purposes of these experiments serves as the paradigmatic medium chain fatty acid. 5mM was chosen as a starting concentration based upon McDonnell et al.^20^ Our initial hypothesis was that lipids were altering histone acetylation rather than methylation as OA-mediated histone acetylation had been demonstrated for both the MCF7 and MDA-MB-231 cell lines in McDonnell et al.^20^ Therefore, histone acetylation was used as a readout to determine concentration. We observed H3K14 acetylation at both 2 mM and 5 mM OA but H3K9 acetylation only at 5 mM^14^; therefore, we continued experiments at 5 mM. Subsequent analysis of histone acetylation and methylation utilizing the Epiproteomic Histone Modification Panel revealed that OA increased histone methylation more significantly than histone acetylation; this fact borne out by subsequent methylation and acetylation flux analysis.^14^

In this study, we performed metabolic and epigenomic studies in MCF-10A cells as well as scRNA-seq studies in tissue-derived breast microstructures to identify the mechanism linking lipid metabolism, epigenetic reprogramming, and the genesis of ER-BC. We demonstrated that the metabolism of OA in preference to glucose and glutamine results in a metabolic shift toward the SOG and methionine pathways increasing the production of SAM, glutathione, and 2-HG which can produce profound consequences on the mammary cells (Fig. 8). Metabolic flux-based calculations showed that OA increased the cancer index which is defined by the Warburg effect. The promotion of flux into SOG and methionine pathways is very probably initiated by OA induced the increased expression of PHGDH and phosphoserine aminotrasnferase (PSAT1), the enzyme catalyzing the second step in the *de novo* serine pathway, which was observed both in MCF-10A cells and tissue derived breast microstructures. ATF3 and ATF4 are both induced by endoplasmic stress (ER stress); one of the causes of ER stress is oxidative stress, which we have demonstrated in the ROS studies (Fig. 2). Both ATF3 and ATF4 are transcriptional regulators of the serine metabolic pathway genes^21,22^; ATF3 expression is increased in almost all cell subtypes in the scRNA-seq study. PHGDH and PSAT1 are upregulated in ER-BC and have an important role in cancer development.^21,23^ PHGDH expression is elevated in approximately 70 % of ER-BCs and the SOG pathway gene signature is significantly correlated with ER-status.^24,25^ PHGDH also produces the oncometabolite 2-HG from α-KG in the thermodynamically favored reverse direction.^26^ The appearance of DNA breaks are likely consequent to 2-HG increase with probably the contribution of the increase of ROS.^27^ Sulkowsi and collaborators showed that the overproduction of 2-HG as a consequence of mutations in isositrate dehydrogenase 1 and 2 impairs DNA repair. 2-HG inhibits the αKG-dependent dioxygenases such as KDM 4A/B. Their catalytic activity is required for homologous recombination repair; inhibition results in metabolic “BRCAness”.^16^

**Fig. 8:**
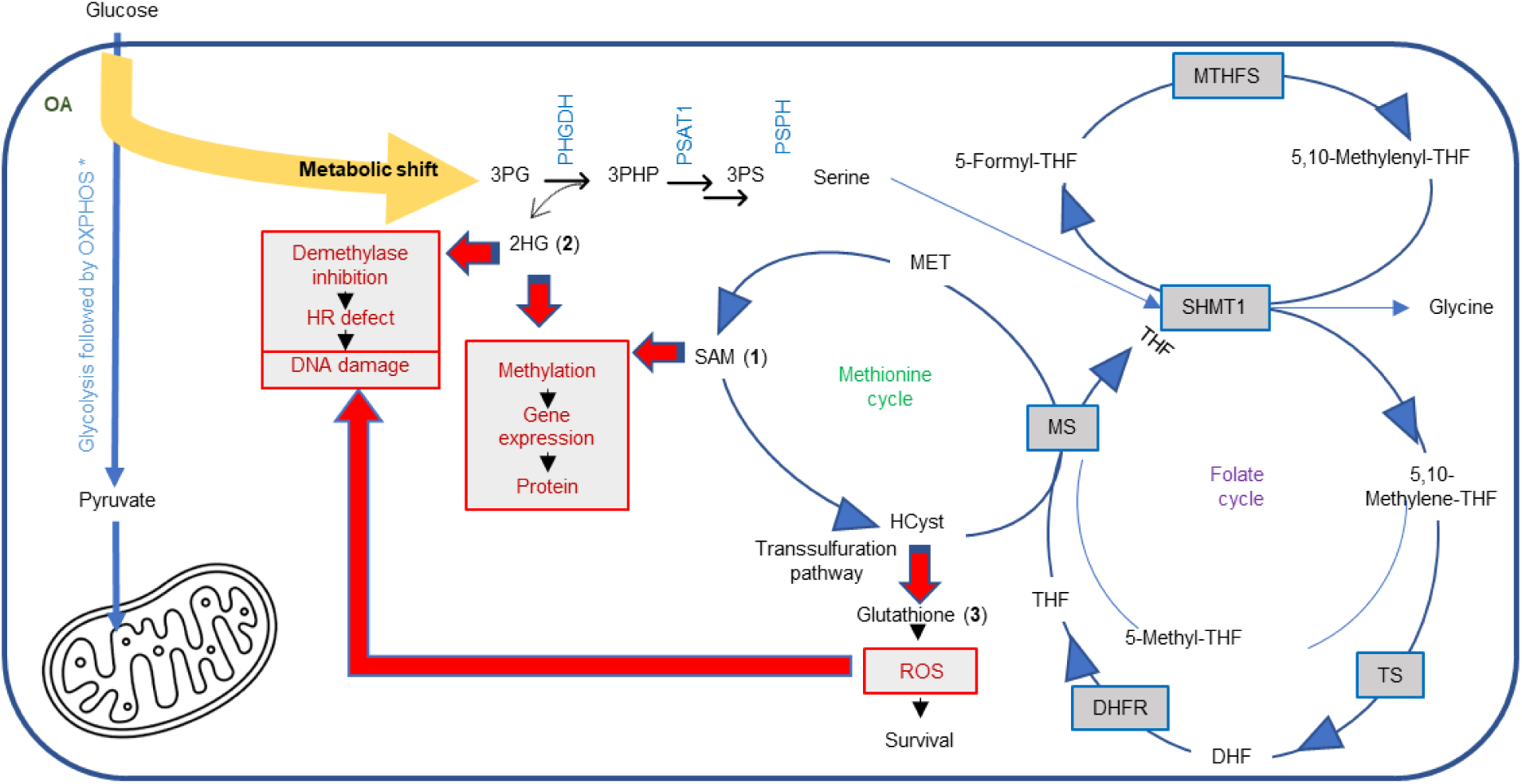
OA induced shift of metabolism away from glycolysis to the SOG/ methionine cycle. Metabolic shift leads to: (1) increased SAM, which is responsible for epigenetic induced plasticity; (2) increased 2-HG, which impairs DNA repair and contribute to epigenetic fostered plasticity; and (3) increased glutathione (GSH) to counter the ROS and thus enabling survival of a subset of cells and contributing to DNA damage. SHMT1, Serine Hydroxymethyltransferase 1; TS, Thymidylate Synthase; DHF, Dihydrofolic acid; DHFR, Dihydrofolate Reductase; MS, Methionine Synthase; MTHFS, Methenyltetrahydrofolate Synthetase; THF, Tetrahydrofolate; MET, Methionine; HCyst, HomoCysteine.

The increased production of SAM, the universal methyl donor, and 2-HG, a demethylase inhibitor, lead to epigenetic, i.e., H3K4 trimethylation, fostered phenotypic plasticity. This plasticity leads to reprogramming and/or differentiation to cells that express Neural, EMT, stem cell and breast cancer related genes. Intriguingly, network analysis of pathways associated with H3K4me3 peaks upon OA exposure revealed several neural related pathways. Pathways that related to the neuronal system were also overrepresented in breast microstructures upon OA: e.g. signaling by NTRKS (Neurotrophic tyrosine receptor kinase) and NGF stimulated transcription which have been associated with ER-BC,^28^ stem cell self-renewal and plasticity; ^29^ MK signaling which has been involved in neurogenesis and cancer progression;^30^ and synthesis of neurotransmitters which can be produced by non-neuronal cells including BC cells.^31^ Of note, BC is one of the malignancies in which a neural differentiation phenotype has been observed, specifically in Triple Negative BC (TNBC), a subtype of ER-BC.^32^ This specific differentiation, i.e. neural, may be a key feature of tumorigenesis, which is hypothesized to represent the culmination of a process of gradual loss of a cell’s original identity and gain of the properties of NSC/Neural Progenitor Cells.^33–35^

The changes of cell state, e.g. to LumHR-major or to LumSec-HLA, the latter characterized by the expression of the chemokine ligand 20 (CCL20)^19^, has been noted to be involved in breast cancer progression, EMT, migration and invasion.^36^

Live monitoring of ROS-induced redox changes revealed a significant ROS increase shortly after OA exposure. ROS are controlled by antioxidant defenses, this probably favors/enables the survival of specific cell subtypes that can mitigate ROS. Metabolic flux analysis of our scRNA-seq data revealed glutathione metabolism and ROS detoxification was increased only in BSL1. This may not be surprising as previous scRNA-seq of primary human breast epithelial cells has revealed that the three subtypes of breast epithelial cells: luminal mature (or HS), LPs and basal, have distinct metabolic properties.^37^ Previous studies have demonstrated that basal cells maintain low levels of reactive oxygen species (ROS) primarily by engaging a GSH-dependent antioxidant mechanism while LPs have a robust antioxidant defense as they can mitigate higher levels of ROS relying on multiple GSH-independent antioxidants;^38^ this may be reflected in the increase survival of LP3 cells despite the downregulation of glutathione metabolism and ROS detoxification. LPs are posited to be the cells of origin of basal-like BCs (an ER-BC subtype) and it has been shown that under non-physiological conditions they can acquire stem-like features suggesting a remarkable degree of plasticity. It is possible that a subset of LPs are the ones that can survive and change facilitating malignant transformation, however further experiments would need to be performed to explore this possibility.

Our data represents a compendium of *in vitro* and ex vivo metabolic, epigenomic and molecular data on the effects of the medium-chain saturated fatty acid OA, in the context of human plasma-like medium. OA freely diffuses across cell membranes due to its small size and lipophilic nature, enabling the delivery of a reliable concentration, and reproducibility across experiments. We have assayed gene expression changes in MCF-10A cells consequent to exposure to the long chain, unsaturated fatty acid (FA), linoleic acid (LA). The number of statistically significant changes in gene expression were considerably fewer in the LA treated cells when compared to the OA treated cells.^14^ This difference, OA versus LA, has also been observed in adult rat cardiac myocytes as well.^39^ Entry of long chain FAs into the mitochondria is strictly regulated by CPT1 whereas the medium chain FAs freely diffuse into the mitochondria, which may be a significant vulnerability given the profound changes in gene expression we have observed. The use of only one fatty acid is a limitation of our study. We recognized that additional fatty acids should continue to be explored; however, the current paucity of information regarding the lipid content of the normal breast limits our ability to make an informed selection. Mammary cells have access to lipids from serum or adipocytes, a major component of breast adipose tissue. Mammary cells have the ability to synthesize medium chain fatty acids, which are a component of milk.^40^ From co-culture studies of breast cancer cell lines and adipocytes, it has been shown that adipocyte-derived fatty acids drive proliferation and migration.^41^ Lipid uptake, for example that mediated by fatty acid translocase/CD36 and Fatty Acid Binding Protein 4 (FAB4) facilitate fatty acid transport into the mammary epithelial cells and have been shown to play critical roles in proliferation, migration and metastasis of breast cancer cells.^42,43^ Although the relationship between dietary fat intake and breast cancer risk remains controversial; breast adipose tissue measurement in pre and postmenopausal women, and breast lipid composition imaging evaluation in postmenopausal women revealed an association between saturated fatty acids and breast cancer.^44,45^ In addition, a high saturated fat diet during puberty or adulthood increased the incidence of mammary tumors in mouse models.^46^ The few studies describing the effects of OA in the context of cancer are mixed. Narayaran et al. proposed that OA may have anticancer properties based on OA antiproliferative effect in skin, colon and breast cancer cell lines; however, we and other have shown OA’s oncogenic properties in normal mammary cells,^14^ bladder cancer, and prostate cancer. ^47,48^

We have demonstrated that exposure to fatty acids can produce profound consequences on mammary cells/tissue. The effects intersect many of the hallmarks of cancer including deregulating cellular metabolism, unlocking phenotypic plasticity^49^ and a recently added new hallmark: the intersection with neurobiology^50^. In discussing this new hallmark, Hanahan and Monje reflect on the fact that the expression of neuronal signaling and regulatory circuits are in observed in cancer cells of multiple origins, not just ones with ontological relationships to neurons.^50^ They also note cancer cells also can exhibit distinctly neuronal structural features, such as the extension of long, neurite-like processes that facilitate cell to-cell communication in the tumor microenvironment.^50^ They also note cancer cells also can exhibit distinctly neuronal structural features, such as the extension of long, neurite-like processes that facilitate cell to-cell communication in the tumor microenvironment. Axonogenesis and neurogenesis are observed in premalignant lesions, for example, in the initial stages of prostatic intraepithelial neoplasia (PIN)^51^, however, the data we have presented here is, to our knowledge, the first demonstration that this occurs in normal tissue and is consequent to lipid exposure. Our findings unlock the possibility of novel preventive strategies; for example blocking the first enzyme of the *de novo serine* pathway to block the OA-induced increase of SAM.

## Methods

### Cell culture

MCF-10A cell line was obtained from American Type Culture Collection (ATCC) and cultured in mammary epithelial cell growth basal medium (MEBM) with single quots supplements and growth factors (Lonza, #CC-4136). Alternatively cells were cultured in Human Plasma-like Medium (HPLM, Gibco, #A4899101) supplemented with H14 media additives.^52^ Cells were regularly screened for absence of mycoplasma contamination with the ATCC Universal Mycoplasma Detection Kit (ATCC, #301012K).

### Glucose 13C tracing

For isotope tracing experiments, cells were seeded in biological triplicate (∼80% confluent) in HPLM plus H14 supplements and treated with 5mM of the medium-chain fatty acid OA (Sigma, #C5038) dissolved in PBS, or vehicle (PBS) for 20-23 hrs. Cells were washed once with RPMI 1640 Medium without glucose (Gibco #11879020) and then incubated in RPMI 1640 Medium without glucose containing 900.8 mg/L D-Glucose-^13^C_6_ (Sigma, #389374), supplemented with H14 media additives,^52^ 700 µM Alanine (Sigma #A7627), 150 µM Cysteine (Sigma # C1276), 40 µM Pyruvate (Gibco# 11360070) and either vehicle or 5mM OA. After 1, 2 of 4 hours metabolites were extracted as described previously ^53^. Briefly, cells were rinsed twice with 5 ml of ice-cold saline solution, 1ml of 80 % methanol (Sigma, #34860) cooled to −80°C was added and cells were scraped on dry ice. Cell lysates were incubated at -80°C for 5 minutes and vortexed for 1 minute at room temperature, after repeating these steps twice cell lysates were incubated at -20°C overnight. Lysates were then vortexed for 30 seconds and insoluble material was pelleted by centrifugation at 20 000 xg for 15 minutes at 4°C. Supernatant was collected for High-Performance Liquid Chromatography and High-Resolution Mass Spectrometry and Tandem Mass Spectrometry (HPLC-MS/MS) analysis. Data was analyzed at 4 hours after 13C-label addition.

### Analysis of 13C tracing data

We constructed a metabolic network model that covers most fluxes in glycolysis, TCA cycle, pentose phosphate pathway, one-carbon metabolism pathway, and amino acids synthesis pathway. To reduce random errors in batch preparation and measurement, mass isotopomer distributions (MIDs) of metabolites in all biological repeats are averaged. These average MID data are fitted with the following procedure: MIDs of all target metabolites are predicted by averaging the MID of the precursors, weighted by the corresponding generating fluxes with the presumed value:

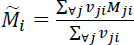

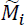: predicted MID vector of metabolite *i; M_ij_*: MID of metabolite *i* produced from a substrate *j*: *v_ij_* the flux from *j* to *i*.

If *M_ji_* is still unknown, it can be deduced by the same procedure, until MIDs of all precursors are known. Then, the difference between the predicted and experimental MIDs of target metabolites was evaluated by Kullback–Leibler divergence:

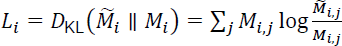

*L_i_* : difference of target metabolite 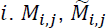: element *j* in vector *M_i_* and 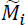 .

Sum of *L_i_* for all target metabolites was regarded as the total difference *L_total_* to minimize by adjusting flux vector v = {*v_i_*} that including all fluxes. Therefore, an optimization problem was defined as:

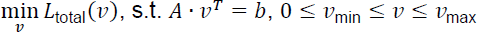

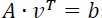: flux balance requirement and other equality constraints; *v_min_*, *v_max_*: lower and upper bounds for composite vector *v*.

The solution of this optimization problem *v\** gives a combination of all fluxes in the network model that fits MID data. Method utilized to solve this optimization problem refers to previous publication^54^. For better precision, accuracy and robustness, the optimization is repeated 10000 times and the 50 solutions with minimal final *L*_total_ are selected to be the final solution set.

### Targeted metabolomics

To determine the relative abundances of intracellular metabolites over time, MCF-10A cells seeded in in biological triplicate in HPLM plus H14 supplements, were treated with vehicle (PBS or PBS plus DMSO), 5mM OA and 5mM OA plus 30uM the PHGDH inhibitor CBR-5884 (Sigma #SML1656). Metabolites were prepared as described in the glucose C13-glucose tracing experiment and collected for LC-MS analysis. The targeted metabolites serine, SAM, glycine, αKG, sarcosine, glycine betaine, methylglyoxal, 2-HG, glutathione, Phosphoribosyl diphosphate (PRPP), betaine aldehyde, and NADH were measured after 0 minutes, 5 minutes, 15 minutes, 1 hour, 2 hours and 4 hours exposure.

### Quantification of D-2-HG and L-2HG

To quantify 2-HG enantiomers D-2-HG and L-2-HG, MCF-10A cells (biological triplicate) seeded in HPLM plus H14 supplements were exposed with vehicle (PBS) and 5mM OA for 30 minutes. In addition, MCF-10A cells were transfected with the human ON-TARGET plus siRNA library targeting PHGDH (Dharmacon) or non-target control (Dharmacon) using the Neon Transfection System (Invitrogen). Then 96 hours post transcription, cells were exposed to exposed with vehicle (PBS) and 5mM OA for 30 minutes. Metabolites were then prepared as described in the glucose C13-glucose tracing experiment. L-2HG and R-2HG derivatization was performed as previously described. ^55^. The amount of L-2HG and R-2HG in extracts was quantified by using a calibration curve and normalized to number of cells.

### Analysis of mitochondrial and nuclear ROS using in vivo reporters

ROS-induced redox changes were monitored using ORP1-roGFP2 based sensors in MCF-10A cells. NLS-roGFP2-Orp1 and mito-roGFP2-Orp1 (puromycin as selection mark) plasmids were obtained from cloning the c-myc sequence or a MTS sequence respectively into the pEIGW roGFP2-ORP1 vector (Addgene #64993). For lentivirus production, HEK293T/17 cells (ATCC) in phenol red free OPTI-MEM (Gibco) were co-transfected with a construct of interest and packaging plasmids Gag-Pol 8.91 (Addgene #187441) and VSV-G (Addgene #8454) using Lipofectamine 3000 (Thermo Fisher Scientific). Cells were incubated at 37°C and after 24h and 48h lentivirus-rich medium was collected, centrifuged (500 x g, 5 min). Supernatant was collected, filtered (0.45 µm) and used to transduction. Transduced MCF-10 cells were cultured in the appropriate medium for 48h and selected with antibiotics. Cells were imaged in Lionheart FX (Biotek). roGFP2-Orp1 was excited sequentially at 405 nm and 488 nm and the emission was recorded at 525 nm. The generated images were analyzed using FIJI ImageJ (1.53c). Each channel was converted to 8-bit and Median (radius = 2 pixels) and Gaussian blur (radius = 2 pixels) were applied. The background was subtracted and a threshold was set to avoid artifacts. Only the nuclear or mitochondrial area were considered for the analysis. Oxidation levels were then determined on a pixel by pixel basis dividing the signal emitted for each channel (405 nm / 488 nm) using Imaging Expressing Parser. Final ratiometric values were obtained using Analyzing particles (size: 10 µm2 - infinity; circularity: 0.50 - 1.00). Obtained values were plotted in GraphPad Prism 10 and a heatmap (Lookup table: Fire) was created for visualization.

### Comet assay

The alkaline comet assay (Abcam #ab238544) for assessing DNA damage in MCF-10A cells was performed according to the manufacturer’s protocol. Slides were viewed with a Lionheart FX (Biotek) using a FITC filter. Images were analyzed by the ImageJ software (National Institutes of Health) using OpenComet v1.3.1.^56^

### CUT&RUN-Seq

For CUT&RUN cells were seeded in biological triplicate (∼80% confluent) in complete MEBM media and exposed to 5mM OA or vehicle (PBS) for 24 hours. About 900’000 cells were processed using the CUTANA kit (Epicypher, #14-1048). Briefly, MCF-10A cells were detached by exposure to 0.25 % trypsin-EDTA (Gibco, #15400054) for 1 minute, then cells were scrapped off the culture plates and centrifuged at 600 × *g* for 3 min. Pellets were washed twice with CUT&RUN wash buffer, and then mixed with Concanavalin A (ConA) beads. Following a 10 min incubation on a magnet rack at RT, the supernatant was discarded. Samples were re-suspended in antibody buffer, and antibodies were added to the beads along with the bound cells as follows, 1:100 H3K4me3 Antibody (Epicypher, #13-0041), 1:50 H3K27me3 antibody (Cell Signaling, #9733) and, 1:100 Rabbit IgG Negative Control Antibody (Epicypher, #13-0042) and incubated overnight at 4°C. Then supernatant was removed, and beads were washed twice with cell permeabilization buffer. Next, pAG-MNase was added to the sample for 10 minutes at RT follow by two washes with cell permeabilization buffer. Then CaCl_2_ was added to a final concentration of 2 mM to activate MNase and initiate chromatin cleavage. Samples were incubated for 2 hours at 4°C. Following incubation, STOP buffer was added and cells cell incubated at 37°C for 10 min to release chromatin fragments. DNA was purified using the CUTANA DNA purification kit. The purified DNA was quantified using Qubit (Invitrogen) as per manufacturer’s instructions. Library preparation was performed using the NEBNext Ultra II Library Prep Kit for Illumina (New England BioLabs, #E7645S) per manufacturer’s instructions with minor modifications. Following adapter ligation, DNA cleanup was performed using 1.0x AMPure XP beads (Beckman Coulter Inc, # A63880). PCR was performed using unique dual index primer pairs (NEBNext Multiplex Oligos for Illumina from New England BioLabs, # E6440S) according the following parameters: 45 s at 98 °C to activate hot-start Q5 polymerase, followed by 15 s at 98 °C, 10 s at 65 °C for a total of 13 cycles, and finally 1 min at 72 °C for final extension. DNA cleanup was performed using 1.0x AMPure beads (Beckman Coulter Inc. #A63881) and the DNA was eluted in TE buffer. DNA quantification was performed using Qubit (Invitrogen), and fragment sizes of individual libraries were analyzed using the High Sensitivity DNA Kit (Bioanalyzer®). Libraries were pooled to a final concentration of 23 nM and sequenced on NovaSeq 6000 (Illumina), 100 bp paired-end reads.

### CUT&RUN data analysis

MACS2 v2.2.7.1 was used to call H3K4me3 narrow peaks and SICER2 was used to call H3K27me3 broad peaks. The differential binding regions were identified using DiffBind v3.9. For H3K4me3 default summit parameter was used. For H3K27me3, summits= 1000 was used. ChIPseeker 1.8.6 was used to annotate the differential peaks identified by DiffBind. HOMER v4.11 was used to scan for the enrichment of motifs using default parameters. CUT&RUN data was compared to published bulk RNA sequencing data.^14^ The Enrichr web was utilized to perform pathways analysis (https://maayanlab.cloud/Enrichr/, accessed on 14 September 2023).

### Culture in Poly-D-Lysine/Laminin plates

MCF-10A, MCF-12A and DCIS.com cells were plated in Poly-D-Lysine/Laminin (PDL/LM) chamber slides (Corning, #354595) treated with vehicle or OA for 24 hrs. Pictures were taken using an EVOS® Digital Microscope.

### Quantitative reverse transcription PCR (RT-qPCR)

MCF-10A cells were exposed to vehicle (PBS or PBS plus DMSO), 5mM OA, 5mM OA plus 30uM CBR-5884 or 5 mM OA plus 160 µM Piribedil for 24 hrs. RNA was then isolated with Qiagen AllPrep DNA/RNA/Protein Mini Kit (Qiagen # 80004). Breast microstructures from 5 independent donors were exposed to vehicle (PBS and PBS plus DMSO), 5mM OA and 5mM OA plus 30uM the PHGDH inhibitor CBR-5884 for 24 hrs. After incubation microstructures were washed with PBS and RNA was isolated with Qiagen AllPrep DNA/RNA/Protein Mini Kit (Qiagen # 80004). cDNA was synthesized using the SuperScript VILO cDNA synthesis kit (ThermoFisher #11755250). Real-time qPCR was performed using Applied biosystem QuantStudio 5 real time PCR System (Thermo Scientific). PrimeTime qPCR probe assays (Supplementary Tables 4 and 5) and PrimeTime Gene Expression Master mix were purchased from Integrated DNA Technology (IDT). For the fold change analysis, the 2^-delta delta CT was calculated as described elsewhere^57^. For this, gene of interest levels were normalized relative to the mean expression of the three control genes UBB, RPLP0 and TBP (delta CT). The difference between treated and untreated gave the delta delta CT.

### Mammary microstructures preparation

Tissues from women admitted for reduction mammoplasty recruited under an approved IRB protocol (NU15B07) were collected. All participants provided written informed consent. Breast tissue to be processed was transferred into a sterile petri dish, chopped into small pieces using scissors and then transferred to sterile 50 ml tubes containing 1 mg/ml collagenase from Clostridium histolyticum (Sigma Aldrich, #C0130) in Kaighn’s Modification media (Gibco, #21127022) supplemented with 0.5% BSA (Sigma, #SLCM0392) and Antibiotic-Antimycotic (Gibco, #15240062). A 0.22 μm filter was used to filter the media containing collagenase. Falcon tubes were sealed with tape and tissue was gently dissociated on a shaker at 100 rpm and 37 °C, overnight (16 h). The next day, microstructures were collected and washed with PBS by centrifugation (114 × *g* for 5 min). Microstructures were resuspended in fresh HPLM media (Gibco, #A4899101) supplemented with H14 media additives plus Antibiotic-Antimycotic (Gibco, #15240096) and added to ultra-Low Attachment Surface six-well plate (Corning, #CLS3471).

### Single cell preparation

Breast microstructures were exposed to OA or vehicle (PBS) for 24 hours. Microstructures were then collected into a 50-ml conical tube by centrifugation (179 × *g* for 5 min) and washed with HBSS without calcium and magnesium (Gibco, #14175095) containing 0.1% BSA (Ambion, # AM2616). After 5 minutes centrifugation (179 × *g* for 5 min), collected microstructures were incubated with pre-warmed (37 °C) TrypLE (Gibco, #12604013) for 15 minutes in a shaker at 100 rpm and 37 °C. Every 5 minutes the dissociation mixture was pipetted gently using wide-orifice p1000 tips. Pre-warmed complete HPLM media containing 0.1% BSA was added to the cell suspension and cells were collected by centrifugation (200× *g* for 5 minutes). Cells were incubated with 2 ml RBC lysis buffer (Invitrogen, # 00433357) for 1 minute at room temperature and pre-warmed complete HPLM media containing 0.1% BSA was added to the cell suspension and cells were collected by centrifugation (200× *g* for 5 minutes). Cells were resuspended in 1 ml complete HPLM media containing 5 mg Dispase II (Sigma, #D4693) and 0.1 mg DNase I (Stemcell Technologies, # 07469) and incubated for 3 minutes in a shaker at 200 rpm and 37 °C. After adding Complete HPLM media, cells were passed 3 times through an 18G needle attached to a 10-ml syringe and filter through a 40 µm cell strainer. Additional 3 washes with cold complete HPLM media were performed (200× *g* for 5 minutes at 4 °C). Single cells were then suspended HBSS containing 0.1% BSA.

### scRNA-seq

For scRNA-seq, the cells were stained with trypan blue to count and determine cell viability. Cells were suspended in PBS/0.04%BSA. About 10,000 individual cells per sample were mixed with Master Mix and loaded a 10x Genomics Chromium Single Cell instrument according to the manufacturer’s instructions. All the procedures were done using Single Cell 3ʹ Reagent Kits v3 following the 10x Genomics user guide #CG000183. Libraries were run on the Illumina NovaSeq per the standard 10x configuration.

### Processing and annotation of scRNA-seq data

Raw sequencing data were processed using CellRanger version 7.0.1. Raw sequencing reads were aligned against the GRCh38 human genome with corresponding gene model from Ensembl database version 109. The digital expression matrix file containing UMIs (unique molecular identifiers) was also generated. scRNA-seq analysis was performed using Seurat version 4.3.0 and R version 4.2.2.^58^ The following filtering criteria was used: nFeature_RNA > 2000 and nFeature_RNA < 10,000; % mitochondrial gene < 20%; and % ribosomal genes < 30%. DoubletFinder package version 2.0.3 was used to remove doublets. The NormalizeData function was used for normalization.^59^ The R package Harmony version 0.1.1 was used for batch correction and integration. ^60^ Unsupervised clustering identified 25 clusters (resolution of 0.8). FindAllMarkers function in Seurat was used to identified differentially expressed genes in each cluster compared to the rest of the cells (p value < 0.05 and log fold chang ≥ 0.25). SingleR package (version 2.0.0)^61^ was utilized for cell annotation according to the Reed et al.^18^ and Kumar et al.^19^ reference datasets.

### Single Cell Pathway Analysis

Pathway analysis was performed using SCPA (1.2.0) with msigdbr v7.5.1 Reactome pathways.^62^ Reactome pathways are provided in Supplemental Table 6.

### Cell-cell communication network analysis

Cell-cell interactions among cell subpopulations were investigated using R package CellChat version 1.6.1.^63^ For every pair of cell types, cell-cell communication interaction were identified.

### Metabolic pathway inference

The Compass algorithm was used for in Silico single-cell flux balance analysis (FBA).^64^ Metabolic pathway inference: The Compass algorithm was used for in Silico single-cell flux balance analysis (FBA). Each cell subset (BSL1, LP3, HS1) was re-normalized using the Seurat NormalizeData function with relative count ‘RC’ normalization and a scale factor of 10,000. Compass was run using standard settings. FBA reaction penalties for each RECON2 metabolic subsystem were negative log transformed and Wilcoxon rank-sum test was used to identify reaction subsystems predicted to be altered with/without OA treatment. Predicted changes in metabolic activity are represented by a Cohen’s d value (mean divided by the pooled standard deviation).

## Data availability

The CUT&RUN and scRNA-seq datasets generated and analyzed during the current study will be deposited in the NCBI Gene Expression Omnibus. Raw dataset of isotope tracing and metabolomics generated during this study are provided in Supplementary Tables 6 and 7, respectively. qPCR raw data can be found in Supplementary Tables 4 and 5.

## Acknowledgements

We thank Robert H. Lurie Comprehensive Cancer Center Metabolomics core for metabolomics services. We would like to thank The Center for Medical Genomics at the Indiana University School of Medicine for CUT&RUN sequencing. We thank the UIC single cell sequencing pilot for scRNA-seq. We are grateful to our many lab colleagues for constructive feedback. Research supported by the 2023 AACR-Pfizer Breast Cancer Research Fellowship, Grant Number 23-40-49-BUST (M.B.E.); the Breast Cancer Research Foundation and Bramsen-Hamill Foundation (S.A.K.).

## Author information

### Contributions

Conceptualization and project design: M.B.E., S.E.C., and S.A.K.; Design and interpretation of metabolomics: N.S.C., Metabolomics experiments: M.B.E., P.G.; Metabolic analysis: M.B.E., S.E.C., S.L., J.L.; CUT&RUN analysis: M.B.E, G.C., H.G., X.X., qPCR: M.B.E., J.S., Analysis of ROS: F.R.P, M.G.B, scRNAseq experiments: M.B.E., P.M.Z., M.V.B., scRNAseq analysis: A.B.M.M.K.I., E.V.B., C.W.M., Metabolic flux analysis: C.W.M., R.K.; writing: M.B.E., S.E.C., and S.A.K.

