## Extended data for "_Metabolic_ shift to serine pathway induced by lipids confers oncogenic properties in non-transformed breast cells"

**Supplementary methods**

**Western blotting**

Cells were seeded and treated the next day for the indicated times. After each treatment, cells were collected, washed with PBS, and lysed in radio immunoprecipitation assay (RIPA) buffer (ThermoFisher Scientific # 89900) including protease inhibitors (ThermoFisher Scientific # 78430). Concentration of proteins was estimated using the BCA protein assay kit (ThermoFisher Scientific # 23227). Samples were then loaded on 4–12% Bis Tris acetate gel using MES buffer, blotted on a polyvinylidene fluoride (PVDF) membrane (Bio-rad, 0.2 µM) and blocked with blocking buffer (5% skimmed milk) for 1 h at room temperature. PHGDH primary antibodies was purchased from Cell Signaling Technologies (rabbit mAb Cell Signaling # 66350) and used at 1:1000 dilution. GAPDH primary antibody (mouse mAb, Meridian Life Science # H86504M) was used at 1:10000 dilution. Membranes were incubated in primary antibody overnight at 4 degrees Celsius with shaking. Blots were washed with TBST, three times 5 min each, and probed with secondary antibodies (Anti-rabbit IgG, HRP-linked Antibody, Cell Signaling #7074; or Anti-mouse IgD, HRP-linked Antibody, Cell Signaling # *34709*) at a concentration of 1:10,000 for 1 h at room temperature.

**Generation of PHGDH overexpressing MCF-10A cells**

MCF-10A cells were transduced with lentiviral vectors prepared and used as previously described [23]. In brief, MCF-10A cells transduced with the lentiviral vector pLvx-Tight-Puro encoding for PHGDH. Stably transduced cells were selected with 1 µg/ml puromycin (Gibco, # A1113803 ). PHGDH was induced with 2 µg/ml doxycycline (Sigma-Aldrich, # D9891).

**ALDEFLUOR® assay**

For ALDEFLUOR® assay MCF-10A cells seeded in HLPM plus H14 additives were exposed to 5mM OA or vehicle (PBS) for 24 hours in biological triplicates. The ALDEFLUOR® assay (Stemcell Technologies, **# 01700**) was used according to the manufacturer's instructions. Briefly, 1 x $10^6$ cells/mL cells were suspended in ALDEFLUOR assay buffer containing the ALDH substrate BODIPYaminoacetaldehyde and incubated at 37°C for 60 min. For each sample, cell aliquots were incubated with or without 50 mM diethylaminobenzaldehyde (DEAB), an ALDH-specific inhibitor. ALDH analysis was performed using a FACSymphony A5-Laser Analyzer.

### Extended data

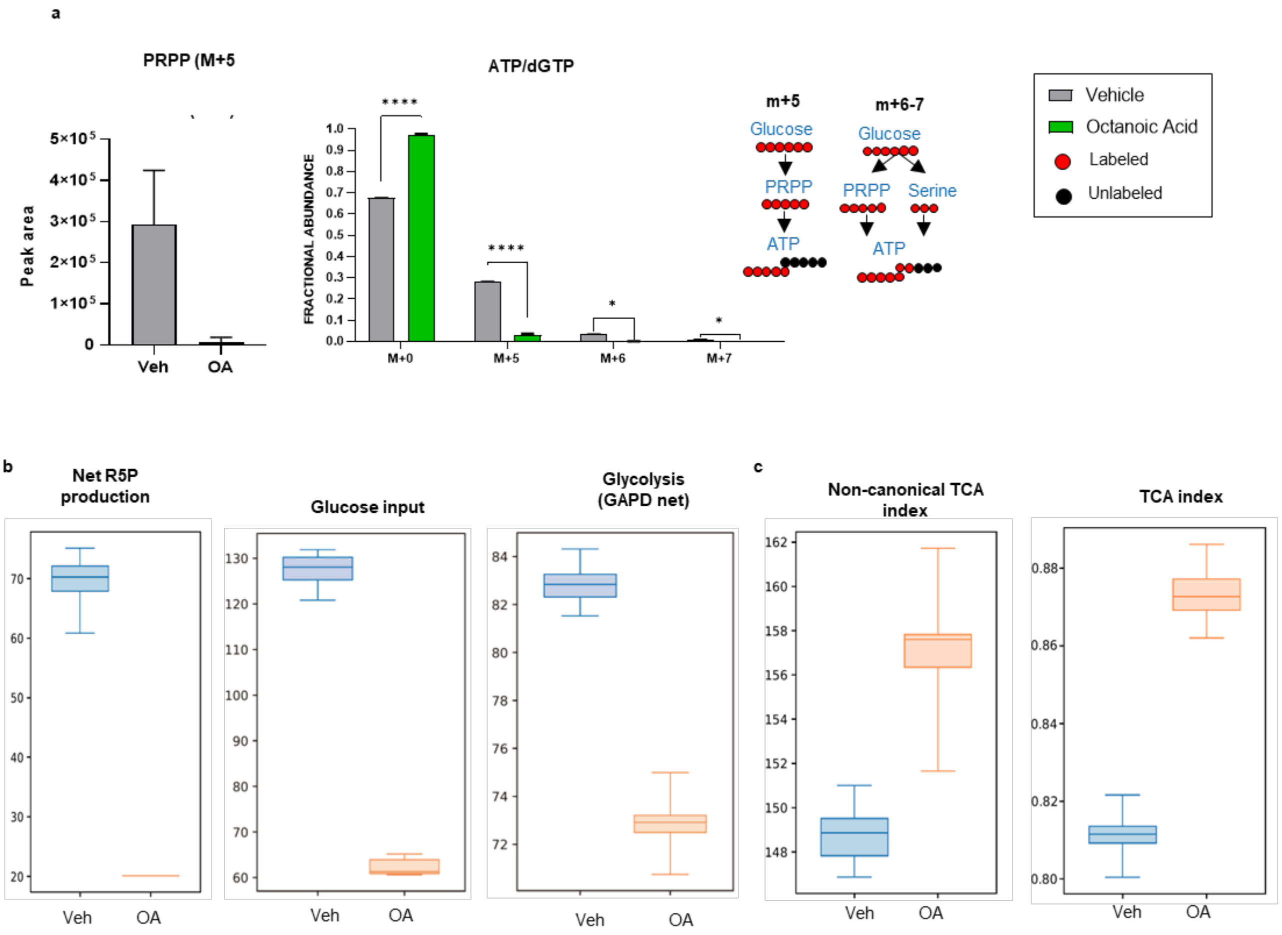


**Extended Data Fig. 1: $^{13}$C metabolic flux analysis**

**a,** Fractional abundance of phosphorilbosyl pyrophosphate (PRPP) M+5 and ATP isotopologues M+5-M+7. *p < 0.05, ****p < 0.0001 (2way anova with Turkey test, n=3, mean with sd). 4 hours labeling. On the right schematic of derivation and contribution of carbon atoms in  ATP synthesis.

**b,** 13C Metabolic flux calculations: Net R5P production, Glucose input, Glycolisis, Non-canonical TCA index and TCA index.

**c,** 13C Metabolic flux calculations: Non-canonical TCA index and TCA index.

a

| Gene | Gene Summary |
|---|---|
| PCDH1 | Involved in neural cell adhesion (possible role in neuronal development). |
| LORICRIN | Major keratinocyte cell envelope protein |
| MIR4652 | |
| NIBAN1 | May be involved in the endoplasmic reticulum stress response. |
| HEY2-AS1 | |
| RAB3C | Member of the RAS oncogene family |
| TNFRSF19 | May play an essential role in embryonic development. |
| LOC105371811 | |
| PLAG1 | Proto-oncogene, stem cell marker |
| SLC30A3 | Zinc ion transporter mediating the import of zinc from cytoplasm into synaptic vesicles |
| C3orf70 | May play a role in neuronal and neurobehavioral development |
| LGR6 | Genome-wide association studies identified LGR6 as an ER-negative and triple-negative-specific breast cancer germline susceptibility gene |

b

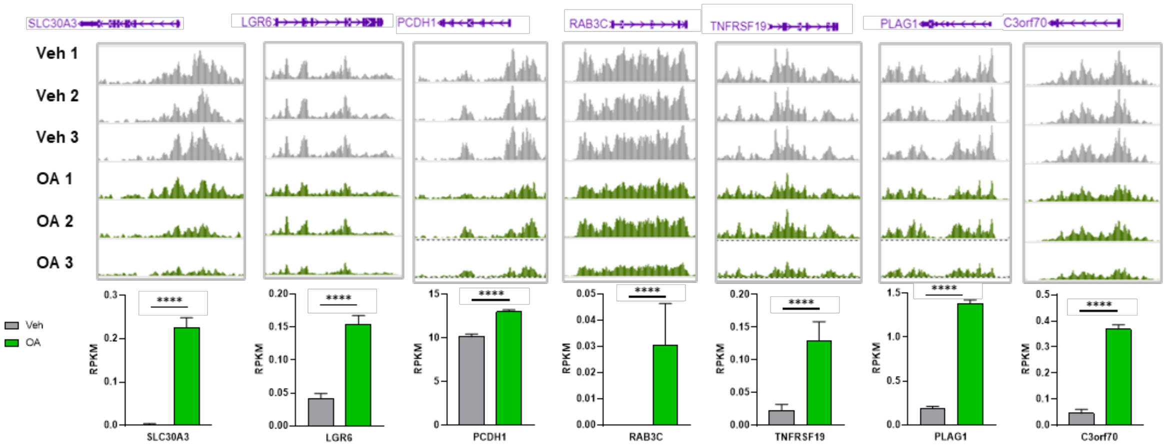


c

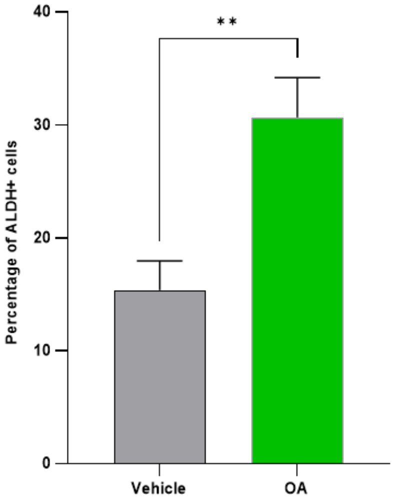


**Extended Data Fig. 2. CUT&RUN for H3K27me3 upon Octanoic acid (OA).**

**a,** Table depicting all genes associated with H3K27me3 peaks. Gene summary is from UniProtKB/Swiss-Prot Summary for each gene.

**b,** Genes differentially modulated by OA with H3k27me3 peaks.

**c,** Identification of stem-like cells (ALDH+) using ALDEFLUOR test in vehicle and OA treated MCF-10A cells.

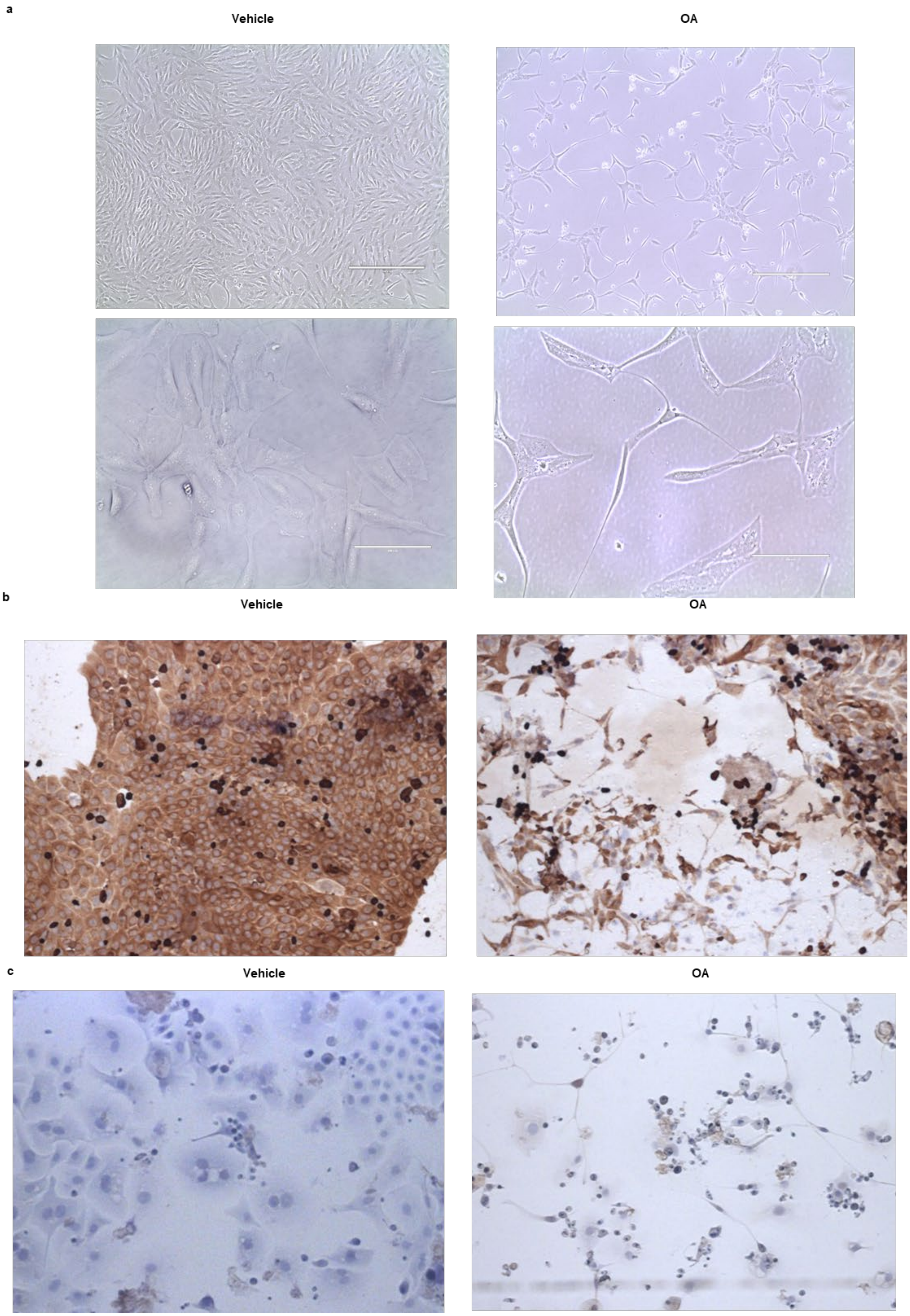
a
Vehicle
OA
b
Vehicle
OA
c
Vehicle
OA

**Extended Data Fig. 3. Breast epithelial cells adopt a neural-like phenotype upon Octanoic acid (OA)**

**a,** MCF-10A cells growing in PDL/L plates following exposure to vehicle (left) or 5 mM OA (right) for 1 week. Magnification: 10x (Upper), 40x (bottom).

**b,** Micrographs of MCF-10A cells growing in PDL/L plates following exposure to vehicle (left) or 5 mM OA (right) for 24 hrs. Magnification: 4x. PanCK and Dapi staining.

**c,** Micrographs of DCIS.com cells following exposure to vehicle (left) or 5 mM OA (right) for 24 hrs. Magnification: 4x. Dapi staining.

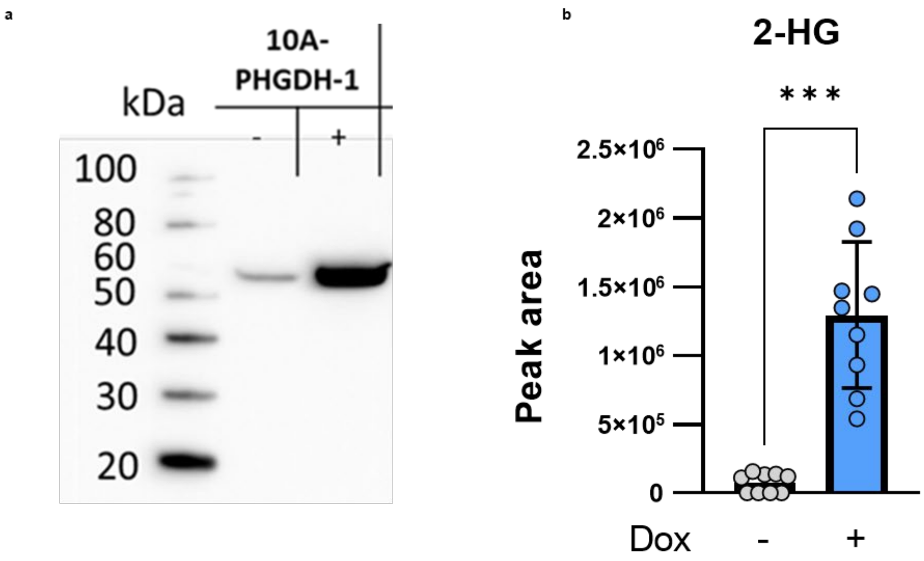


**Extended Data Fig. 4. 2-HG**

**a,** Protein expression of PHGDH by protein blot analysis of 2-HG in MCF-10A cells expressing doxycycline induced PHGDH after 24 hrs exposure to ± 2 µg/ml doxycycline (Dox).

**b,** Measument of 2-HG in MCF-10A cells expressing doxycycline induced PHGDH after 24 hrs exposure to ± 2 µg/ml doxycycline (Dox).

a

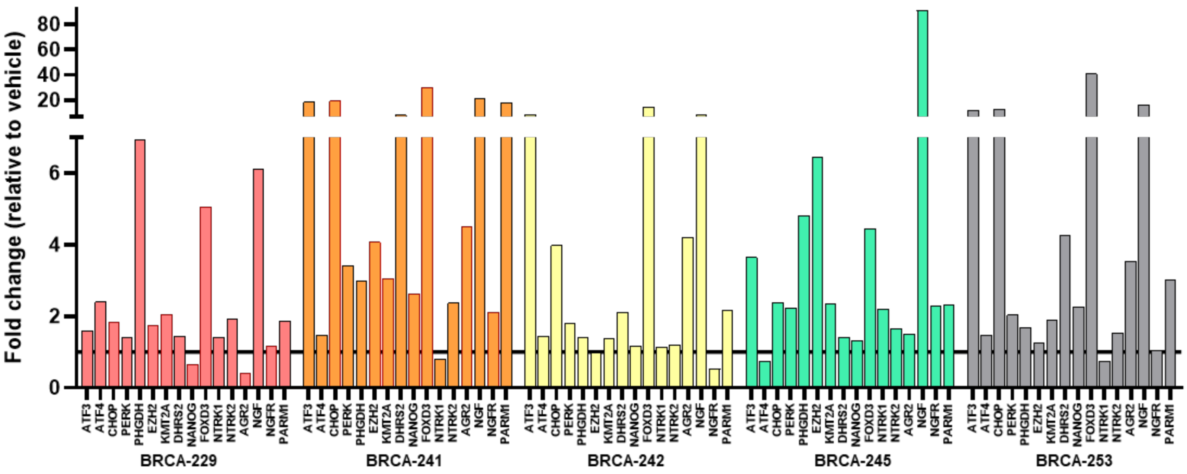


b

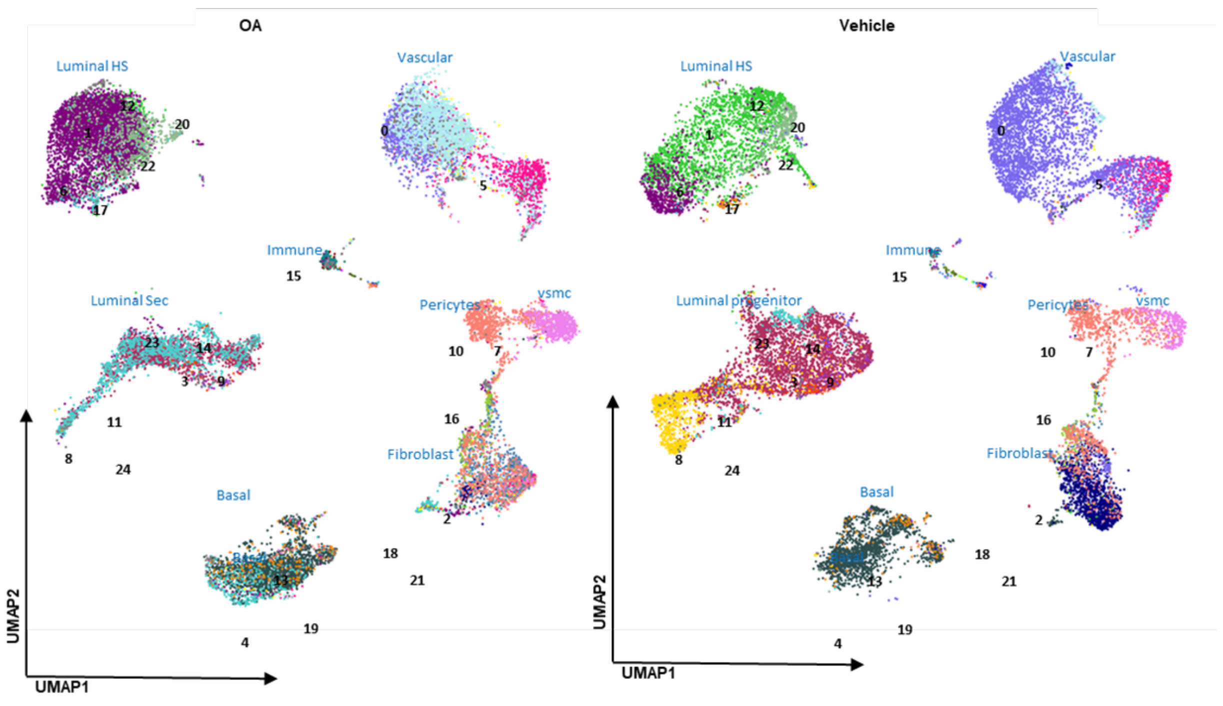


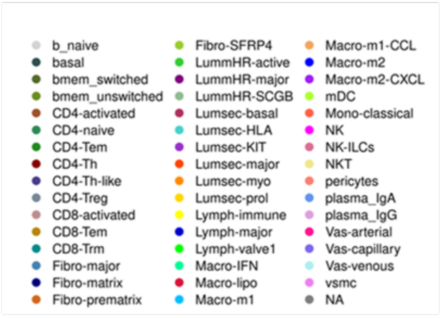


| | %Veh | %OA | | %Veh | %OA | | %Veh | %OA |
|---|---|---|---|---|---|---|---|---|
| basal | 10.5 | 11.2 | b_naive | 0.01 | 0.00 | Macro-lipo | 0.07 | 0.01 |
| LummHR-active | 13.5 | 6.1 | bmem_switched | 0.06 | 0.04 | Macro-m1 | 0.05 | 0.02 |
| LummHR-major | 3.5 | 20.3 | bmem_unswitched | 0.06 | 0.01 | Macro-m1-CCL | 0.02 | 0.08 |
| LummHR-SCGB | 2.1 | 3.2 | CD4-activated | 0.03 | 0.00 | Macro-m2 | 0.11 | 0.04 |
| Lumsec-basal | 19.4 | 8.5 | CD4-naive | 0.03 | 0.40 | Macro-m2-CXCL | 0.00 | 0.02 |
| Lumsec-HLA | 1.1 | 9.4 | CD4-Tem | 0.00 | 0.01 | mDC | 0.04 | 0.00 |
| Lumsec-KIT | 0.5 | 0.2 | CD4-Th | 0.06 | 0.23 | Mono-classical | 0.03 | 0.06 |
| Lumsec-major | 0.5 | 0.1 | CD4-Th-like | 0.04 | 0.21 | NK | 0.00 | 0.01 |
| Lumsec-myo | 1.0 | 0.8 | CD4-Treg | 0.02 | 0.04 | NK-ILCs | 0.00 | 0.01 |
| Lumsec-prol | 5.1 | 0.0 | CD8-activated | 0.01 | 0.01 | NKT | 0.00 | 0.01 |
| Fibro-major | 0.7 | 4.5 | CD8-Tem | 0.00 | 0.01 | plasma_IgA | 0.08 | 0.18 |
| Fibro-matrix | 7.6 | 0.4 | CD8-Trm | 0.06 | 0.30 | plasma_IgG | 0.01 | 0.03 |
| Fibro-prematrix | 0.0 | 0.0 | Lymph-immune | 0.14 | 1.21 | Vas-arterial | 3.32 | 5.54 |
| Fibro-SFRP4 | 0.7 | 0.9 | Lymph-major | 0.14 | 0.20 | Vas-capillary | 20.90 | 4.65 |
| pericytes | 6.2 | 8.2 | Lymph-valve1 | 0.01 | 0.01 | Vas-venous | 0.89 | 9.63 |
| | | | Macro-IFN | 0.01 | 0.01 | vsmc | 1.60 | 3.24 |

**Extended Data Fig. 5: Gene expression analysis of tissue derived breast microstructures.**

**a,** Boxplots showing the expression of OA-induced genes upon OA in tissue-derived breast microstructures from 5 donors measured by qPCR.

**b,** Approximation and Projection (UMAP) plot of 36’904 cells identified a total of 25 different cell clusters. Cell subtypes defined by Kumar et al. Table shows cell subtypes proportions.

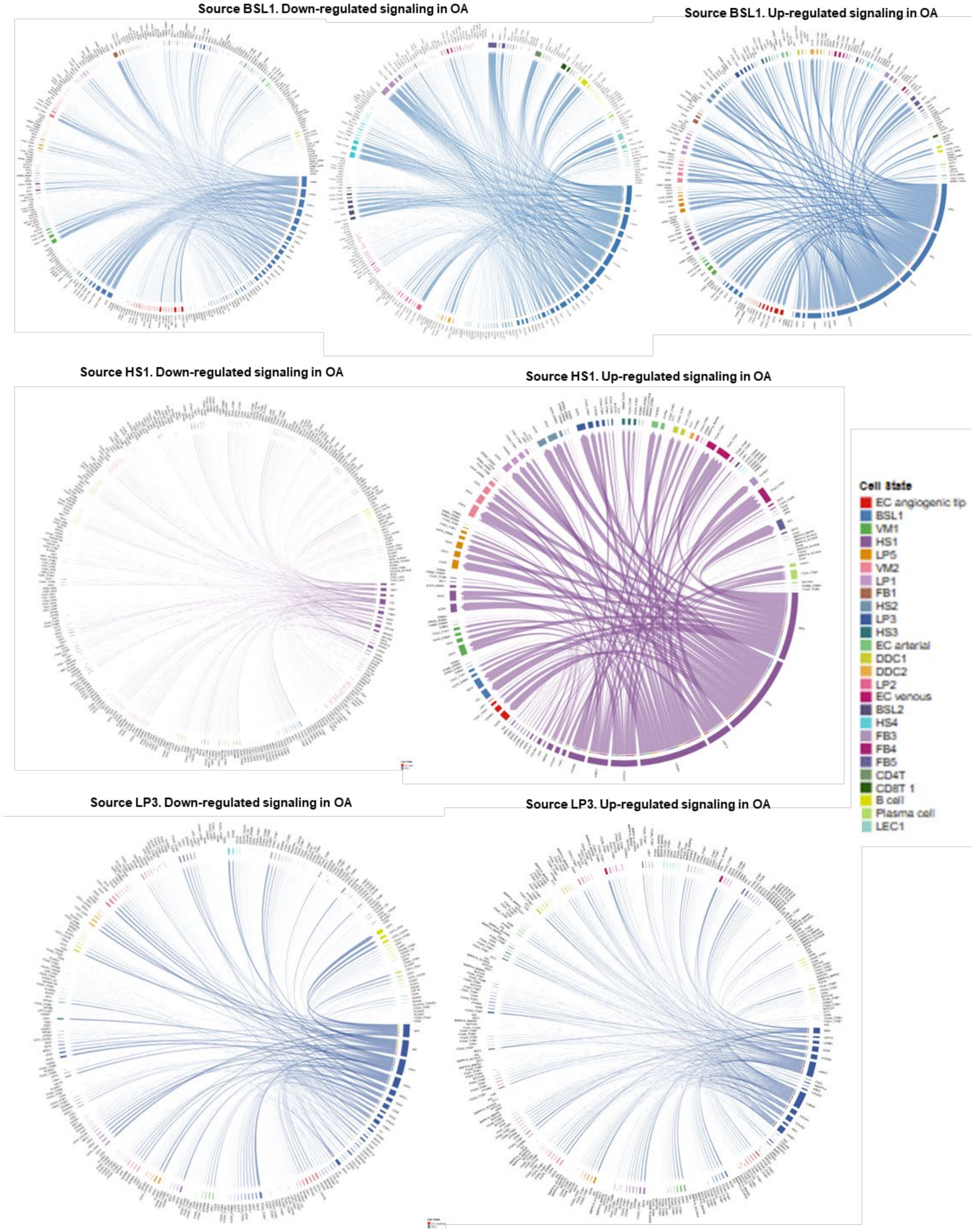


**Extended Data Fig. 6: Major signaling pathways in breast microstructures upon octanoic acid (OA) exposure.**

Chord diagram showing each ligand-receptor pattern and their weights in the interaction between BSL1, HS1, LP3 and all cell subtypes.
